# Droplet Hi-C for Fast and Scalable Profiling of Chromatin Architecture in Single Cells

**DOI:** 10.1101/2024.04.18.590148

**Authors:** Lei Chang, Yang Xie, Brett Taylor, Zhaoning Wang, Jiachen Sun, Tuyet R. Tan, Rafael Bejar, Clark C. Chen, Frank B. Furnari, Ming Hu, Bing Ren

**Author notes:** Corresponding author: Bing Ren. These authors contributed equally: Lei Chang and Yang Xie.

## Abstract

Comprehensive analysis of chromatin architecture is crucial for understanding the gene regulatory programs during development and in disease pathogenesis, yet current methods often inadequately address the unique challenges presented by analysis of heterogeneous tissue samples. Here, we introduce Droplet Hi-C, which employs a commercial microfluidic device for high-throughput, single-cell chromatin conformation profiling in droplets. Using Droplet Hi-C, we mapped the chromatin architecture at single-cell resolution from the mouse cortex and analyzed gene regulatory programs in major cortical cell types. Additionally, we used this technique to detect copy number variation (CNV), structural variations (SVs) and extrachromosomal DNA (ecDNA) in cancer cells, revealing clonal dynamics and other oncogenic events during treatment. We further refined this technique to allow for joint profiling of chromatin architecture and transcriptome in single cells, facilitating a more comprehensive exploration of the links between chromatin architecture and gene expression in both normal tissues and tumors. Thus, Droplet Hi-C not only addresses critical gaps in chromatin analysis of heterogeneous tissues but also emerges as a versatile tool enhancing our understanding of gene regulation in health and disease.

## Introduction

In eukaryotic cells, chromosomes are organized within the nucleus in a non-random manner^1–4^. Each chromosome is folded into a complex and dynamic structure, forming chromatin loops^5^, topologically associating domains^6, 7^, and A/B compartments^8–10^. Chromatin organization plays a crucial role in both embryonic development^11, 12^ and the progression of various diseases^13,14^. During development, the chromatin structure influences which genes are turned on or off by the distal regulatory elements^15–18^. Abnormal chromatin organization can lead to aberrant gene expression or silencing, contributing to cancer development and progression^14^. Indeed, components of the nuclear machinery that regulates chromatin organization are among the most frequently mutated genes in human cancers^19–22^. Thus, comprehensive analysis of the chromatin organization is crucial for understanding the gene regulatory programs involved in both normal development and disease pathogenesis.

Analyzing chromatin organization in primary tissues and tumor biopsies presents unique challenges due to the complexity of chromatin dynamics and the technical limitations of current methodologies. First, the biospecimens available for assays are often heterogeneous, containing a mixture of different cell types. Existing techniques such as *in situ* Hi-C for analysis of 3D chromatin organization typically require bulk tissues as input and could not resolve specific chromatin organization patterns relevant to particular cell types, especially in tumors where cancer cells coexist with stromal and immune cells. Second, the resulting data obtained from bulk chromatin organization assays are complex, and relating changes in chromatin organization to specific functional outcomes can be difficult. Addressing these challenges requires the development of more sensitive single-cell chromatin analysis, better methods for handling tissue samples, and advanced computational tools for data analysis.

Great strides have been made in recent years in single-cell Hi-C technologies^1–4^. In general, single-cell Hi-C methods can be categorized into low-throughput microwell-based approaches and high-throughput combinatorial indexing-based approaches. In the case of microwell-based single-cell Hi-C methods, cells/nuclei are individually dispensed into microwells, and library construction is carried out in each microwell in parallel, frequently with the help of automated liquid handlers. Notably, several single-cell Hi-C methods have been developed, such as those by Nagano *et al*. (2013)^23^, Nagano *et al*. (2017)^24^, Stevens *et al*. (2017)^25^, Flyamer *et al*. (2017)^26^, Tan *et al*. (2018)^27^ and third-generation sequencing (TGS) based scNanoHi-C^28^. Recently, additional microwell-based single-cell Hi-C methods have emerged that combine Hi-C analysis with assays for one or more additional molecular modalities in a single cell. For example, Methyl-HiC^29^ and single-nucleus methyl-3C sequencing (sn-m3C-seq)^30^ capture chromatin interactions and DNA methylation patterns from the same cell. HiRES^31^ jointly performs Hi-C and RNA sequencing to explore the functional relationship between 3D genome organization and transcriptome dynamics. In general, microwell-based techniques are limited in scale, and difficult to adopt broadly due to lengthy procedures and high cost. On the other hand, combinatorial indexing-based single-cell Hi-C methods utilize combinatorial barcoding strategies to achieve high throughput and scalability. Previously, a single-cell combinatorial indexed Hi-C (sciHi-C)^32^ method allowed the generation of single-cell Hi-C libraries from a few thousand cells, albeit with limited genomic coverage in each cell. Building upon the same strategy, GAGE-seq (genome architecture and gene expression by sequencing)^33^ was developed to simultaneously profile chromatin interactions and gene expression in single cells, to achieve high throughput, multimodality, and high coverage per cell. However, the lengthy and largely manual combinatorial indexing procedure still poses a challenge for its general adoption.

Here, we introduce a highly scalable, generally accessible, droplet-based single-cell Hi-C method, Droplet Hi-C, which combines an *in situ* chromosomal conformation capture (3C) assay with commercially available droplet microfluidics, to simultaneously capture the 3D genome structure from tens of thousands of individual cells in a single experiment. We demonstrate the utility of Droplet Hi-C data in resolving cell type-specific chromatin architecture in complex tissues such as the mouse brain. We further show that Droplet Hi-C can be used to identify aberrant chromatin structure in cancer cells. Most excitingly, we used Droplet Hi-C to identify extra-chromosomal DNA (ecDNA) and map their chromatin interactome in tumor cells at single-cell resolution. Finally, we extended this method to enable the simultaneous capture of transcriptome and chromatin architecture in single cells.

## Results

### Development of Droplet Hi-C

Droplet Hi-C builds upon the *in situ* Hi-C procedure^5^. It captures spatial proximity genome-wide between chromatin fibers in formaldehyde-crosslinked cells/nuclei through restriction digestion and ligation *in situ* (Fig. 1a). After SDS treatment to remove histone proteins, DNA fragmentation and capture is then carried out in a commercially available microfluidic platform (i.e. 10x Genomics single cell ATAC kit), where cell-specific DNA barcodes are added to the DNA fragments. This is followed by sequencing library construction and next-generation sequencing (NGS) (Fig. 1a). The whole procedure lasts about 10 hours from fixed cells/nuclei to final sequencing libraries, and 8 samples can be processed in parallel, enabling profiling of 40,000 or more cells simultaneously (Fig. 1a-c). The throughput of Droplet Hi-C surpasses plate-based single-cell Hi-C methods by an order of magnitude, offering the shortest experimental duration and hands-on time (Fig. 1b). Given the widespread use of commercial microfluidic systems, ease of the experimental procedure, and relatively low cost (Fig. 1c), Droplet Hi-C has the potential to be rapidly adopted by the research community.

**Fig. 1:**
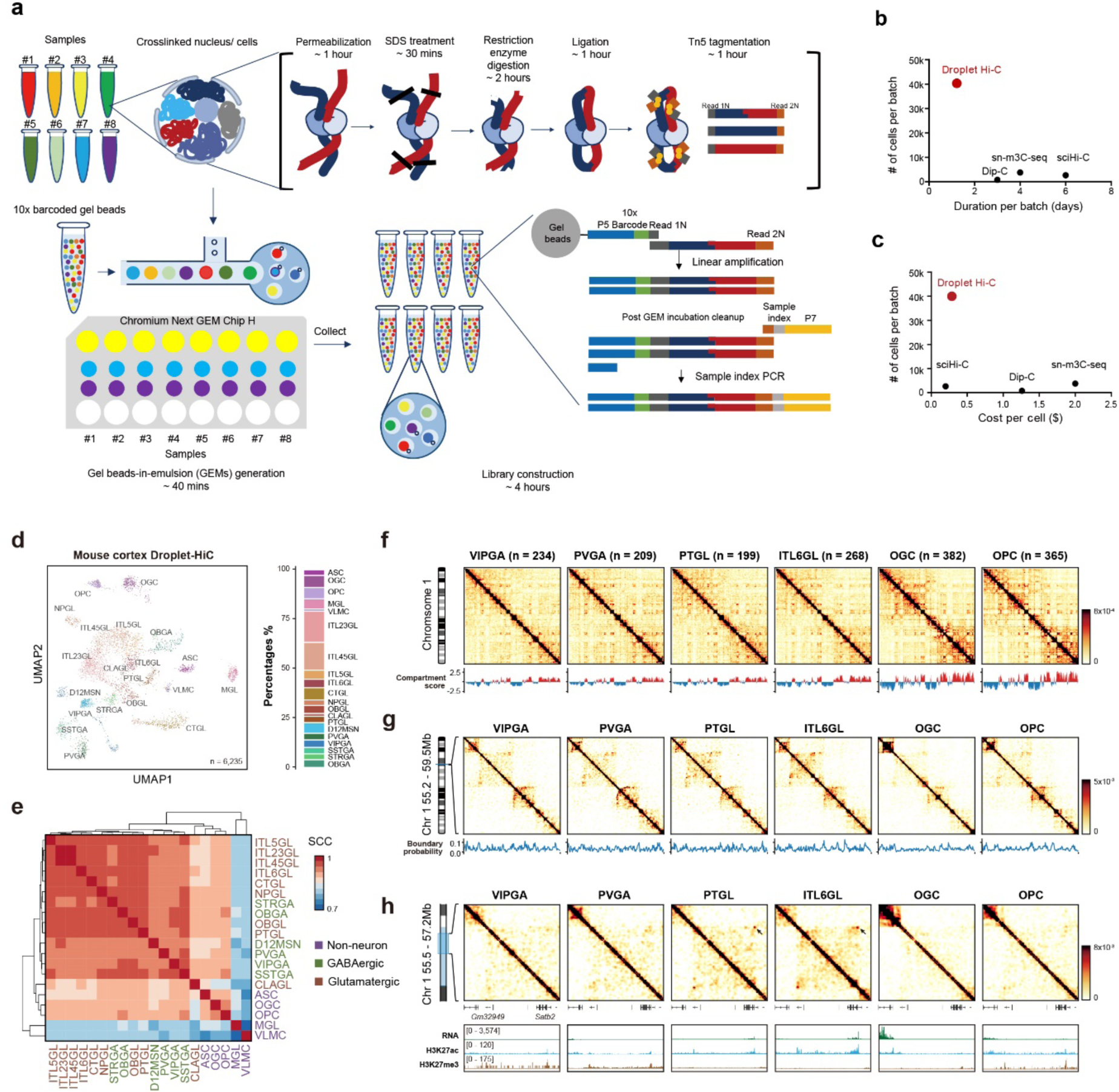
Overview and performance of Droplet Hi-C. **a,** Schematic of the Droplet Hi-C workflow. **b-c,** Comparison of throughput, sample preparation time, and cost among different single-cell Hi-C methods. **d,** UMAP visualization of Droplet Hi-C data from adult mouse cortex. **e,** Genome-wide Spearman’s correlation coefficients (SCC) between compartment scores from different cell types. **f,** Cell type-specific Droplet Hi-C contacts map from chromosome 1 and compartment score at 100-kb resolution. The color bar represents the imputed contact number. **g,** Cell type-specific Droplet Hi-C contacts map and boundary probability of example region (chromosome 1: 55.2 - 59.5Mb) at 25-kb resolution. **h,** Cell type-specific Droplet Hi-C contacts map surrounding gene *Satb2* (chr1: 55.5 - 57.2Mb) at 10-kb resolution, along with genome browser view showing transcriptome and histone modifications profiles in the same cell types from the public datasets.

To test the performance of Droplet Hi-C, we first applied it to a mixture of human HeLa S3 cell line and mouse embryonic stem cell (mESC) line. Equal number of cells from two cell lines were combined after cross-linking and subjected to Droplet Hi-C. The results showed that 1,773 human and 3,489 mouse high-quality cells (with total contacts number > 1,000) were recovered after shallow sequencing, along with 284 potential doublets (Extended Data Fig. 1a). We further carried out Droplet Hi-C with a mixture of three human cell lines (K562, GM12878 and HeLa S3). After filtering, we obtained 3,668 high-quality cells with a median of 73,813 unique contacts in each cell (duplication rate 19.0%), including 1,780 HeLa cells (median 66,828 contacts), 1,209 GM12878 cells (median 76,965 contacts) and also 679 K562 cells (median 77,415 contacts). Using Higashi^34^ for cell clustering, three distinct populations can be clearly distinguished (Extended Data Fig. 1b, Supplementary Table 1). By performing genome wide correlation analysis with bulk *in-situ* Hi-C references, each population can be confidentially assigned to one cell type (Extended Data Fig. 1c-h). On the chromosome level, cell type-specific chromatin structures can be observed, such as the distinct Hi-C pattern of chromosomal rearrangements on chromosome 11 in HeLa S3 cells (Extended Data Fig. 1e). When focusing on a region encompassing the K562-specific upregulated gene *LGR4*^35^, we observed an active A compartment at the *LGR4* genomic locus that is specifically shown in the K562 cells (Extended Data Fig. 1g). The differences in compartment scores as well as insulation scores among different cell lines can also be faithfully reproduced as in the published datasets (Extended Data Fig. 1g-h). In summary, these results demonstrate that Droplet Hi-C can accurately distinguish cell type-specific chromatin organization.

### Droplet Hi-C reveals cell type-specific chromatin structure in mouse cortex

To demonstrate the feasibility of applying Droplet Hi-C to primary heterogeneous tissues, we performed this assay using the cortex dissected from 8-week-old mice. In total, we generated 6,235 high-quality single-cell profiles of chromatin architecture from two replicates, with a median of 175,021 unique contacts per cell (with 22,456 *cis*-long contacts (> 1 kb) and 9,954 trans contacts per cell, at 58% duplication rate) (Extended Data Fig. 2a-c). On the single-cell level, despite the sparse contact signals, we can still discern an enrichment of intra-chromosomal interactions (Extended Data Fig. 2d). To benchmark the performance of Droplet Hi-C, we compared our data with previous single-cell Hi-C methods, Dip-C^36^ and sn-m3C-seq^37^, data from the mouse cortex. The decay curve of contact probability by genomic distance showed a similar trend among these assays, with sn-m3C-seq profiles showing a slightly lower fraction of extreme long-range interactions than Droplet Hi-C and Dip-C (Extended Data Fig. 2e). Droplet Hi-C yields a higher fraction of *cis*-long (> 1 kb) interactions than Dip-C, and lower than sn-m3C-seq (Extended Data Fig. 2f). Among these methods, similar *cis* long-*trans* ratios in chromatin contacts were detected (Extended Data Fig. 2f).

To verify the fidelity of Droplet Hi-C profiles, we examined chromatin structures on contact heatmaps spanning multiple resolutions, and found them to be nearly identical between Droplet Hi-C and sn-m3C-seq (Extended Data Fig. 2g). At the chromatin compartment level, although Droplet Hi-C used Tn5 transposase for chromatin fragmentation in nucleus, bias in interactions within the active A or inactive B compartments is not observed (Extended Data Fig. 2h). The compartment scores and insulation scores from Droplet Hi-C data are highly correlated with other two single-cell Hi-C methods (Extended Data Fig. 2i-l). Overall, these results indicated that Droplet Hi-C can reliably and robustly detect chromatin features in complex tissue.

We leveraged single-nuclei RNA-seq data from previous Droplet Paired-Tag^38^ on mouse cortex to co-embed and annotate our Droplet Hi-C data (Extended Data Fig. 3a, Supplementary Table 2). We successfully resolved 20 cell types, including 5 non-neuronal types, 9 glutamatergic neuronal types and 6 GABAergic neuronal types (Fig. 1d). The cell type prediction results show an overall high prediction score (Extended Data Fig. 3b-c). The cell type identities were confirmed based on concordant annotations when compared to the published cortex sn-m3C-seq data (Extended Data Fig. 3d-e). Among these cell types, neuronal cells exhibit greater similarity to each other than to non-neuronal cells (Fig. 1e). The hierarchical chromatin organization features, such as compartments, domains, and chromatin loops, are distinctly discernible in the aggregated single-cell profiles across various cell types (Fig. 1f-h). For example, the glutamatergic neurons marker gene *Satb2* displayed long-range interactions with the region nearby *Gm32949* gene that are specific in glutamatergic neurons (ITL6GL, Cortex L6 excitatory neurons; PTGL, Cortex NP excitatory neurons), but not in GABAergic (VIPGA, MGE-derived neurogliaform cells *Vip*+; PVGA, MGE-derived neurogliaform cells *Pvalb*+) or non-neuronal cells (OGC, oligodendrocytes; OPC, oligodendrocytes precursor cells) (Fig. 1h). Accompanying with this glutamatergic neurons-specific contact, the *Satb2* gene also displayed an elevated expression level as well as association with an active histone modification, H3K27ac, in ITL6GL and PTGL cells when analyzed in conjunction with our Droplet Paired-Tag data (Fig. 1h). By contrast, in other cell types, this gene remained inactive and was even silenced with the presence of repressive H3K27me3 signals.

To systematically probe the differences in 3D genome structure across different cell types and elucidate their relationships with chromatin state and gene expression, we first examined differences in A/B compartments at 100-kb resolution among cell types. These compartments were found to correspond with transcriptionally active and inactive chromatin states. In total, we identified 895 genomic regions displaying variable compartment scores across the brain cell types (Fig. 2a). When combined with single-cell histone modification profiles previously reported for the mouse cortical region, the compartment scores displayed a positive correlation with H3K27ac signals and a negative correlation with H3K27me3 signals across different cell types, suggesting that variable compartments are associated with dynamic chromatin states (Fig. 2b).

**Fig. 2:**
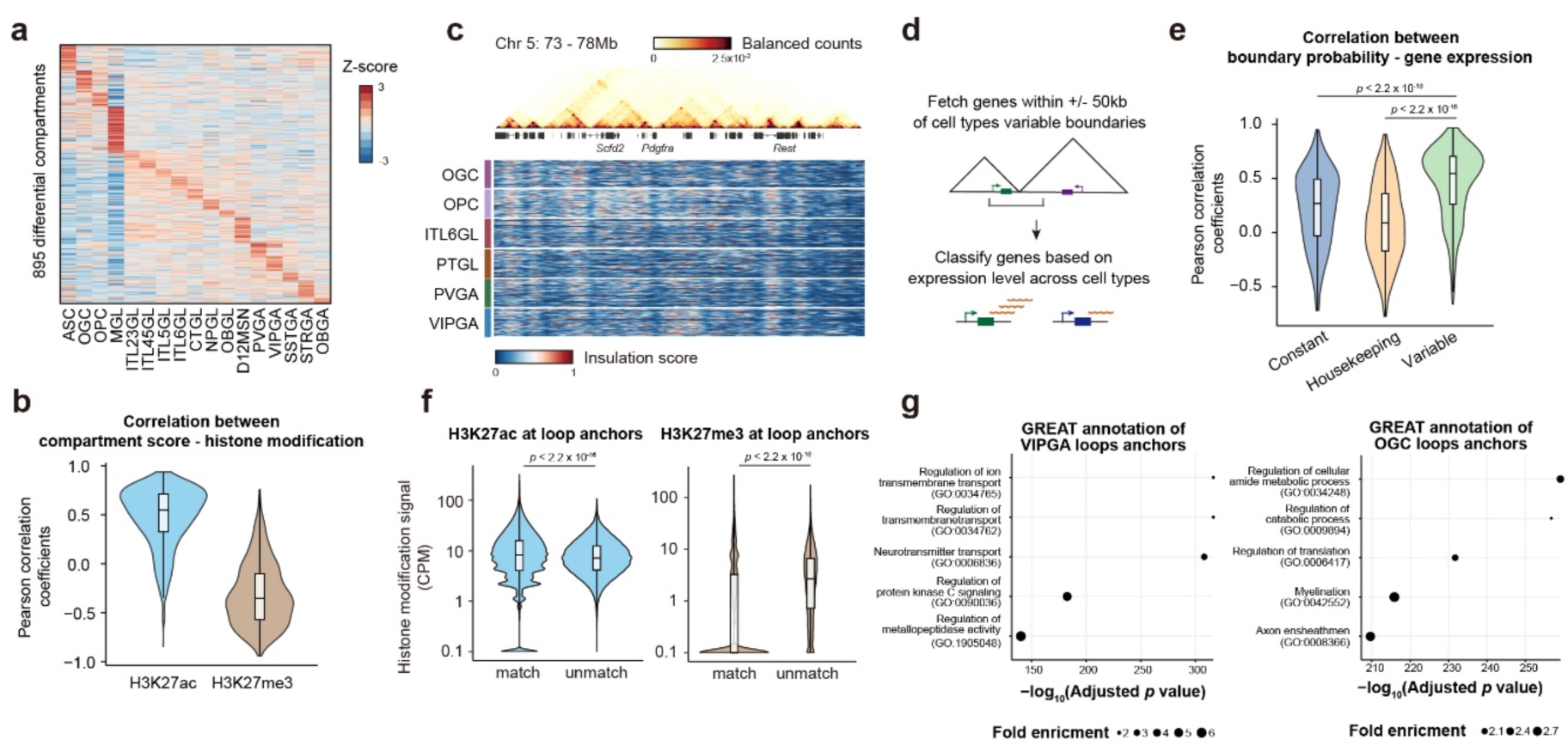
Multi-scale chromatin structures are correlated with cell type-specific gene activity in adult mouse cortex. **a,** Heatmap showing compartment score of differential compartments among all the cell types. **b,** Distribution of Pearson correlation coefficients between compartment score and histone modification signals at each 100-kb bin across differential compartments; n = 895 (H3K27ac and H3K27me3). All box plot hinges were drawn from the 25th to 75th percentiles, with the middle line denoting the median and whiskers denoting 2× the interquartile range. **c,** Comparison of single-cell insulation scores from example cell types surrounding gene *Pdgfra*. A bulk contacts map at 10-kb resolution is shown above. **d,** Schematics for correlation analysis between boundary probability and nearby gene expression level. **e,** Comparison of Pearson correlation coefficients between gene expression level and boundary probability at variable domain boundaries. Genes are classified as constant (n = 521), housekeeping (n = 512) and variable (n = 448) genes. *P* values are from Wilcoxon signed-rank test. All box plot hinges were drawn from the 25th to 75th percentiles, with the middle line denoting the median and whiskers denoting 2× the interquartile range. **f,** Comparison of histone modification signal enrichment at loop anchors across different cell types; n =265,277 (all groups); All box plot hinges were drawn from the 25th to 75th percentiles, with the middle line denoting the median and whiskers denoting 2× the interquartile range. *P* values, Wilcoxon signed-rank test. **g,** Top enriched gene ontology terms for genes at loop anchors in selected cell types VIPGA and OGC. *P* values and fold enrichment were calculated by binomial test. Benjamini–Hochberg FDRs were then calculated to select overrepresented GO terms.

Chromatin domains orchestrate enhancer-promoter interactions and regulate cell type-specific gene activities^18, 36^. We defined domain boundaries in each of the 20 cortical cell types, and subsequently summarized the boundaries across all the cells to obtain the boundary probability for each cell type. Notably, genes with highest expression variation among cell types were often found to reside within or near cell type-specific domain boundaries^34^. For example, for the *Pdgfra* gene, which plays a crucial role in controlling oligodendrocyte differentiation^39^, we found an OPC specific-chromatin domain boundary overlapping the gene body of *Pdgfra* (Fig. 2c). We identified and fetched genes within a 100-kb window centering on variable domain boundaries, and then checked corresponding single-nuclei RNA-seq datasets to classify such genes as constant genes, housekeeping genes and variable genes based on their expression levels across different cell types (Fig. 2d). The results showed that the genes with variable expression levels had higher positive correlation with boundary probability than the constant genes and housekeeping genes (Fig. 2e).

Finally, we examined chromatin loops within each cell type at 10-kb resolution. To check whether loops are associated with functional chromatin states, we compared both H3K27ac and H3K27me3 histone modification signals at loop anchors across all the cell types. Notably, the active chromatin mark H3K27ac showed a higher enrichment at loop anchors identified in the corresponding cell types than other cell types, while repressive chromatin mark H3K27me3 showed the opposite trend, and was more likely to be depleted at the corresponding cell type-specific loop anchors (Fig. 2f). To systematically assess the functions of genes located near cell type-specific loop anchors, we performed GREAT analysis^40^ for VIPGA- and OGC-specific loop anchors. The VIPGA-specific loop anchors are near genes related to neurotransmitters and ion transport, consistent with the signal transduction functions in neurons, while OGC-specific loop anchors are near genes related to myelination and axon ensheathment (Fig. 2g). Taken together, the above analyses showed that Droplet Hi-C can accurately reveal cell type-specific chromatin organization in heterogeneous tissues.

### Droplet Hi-C detects structural alterations and ecDNA in cancer cells

Cancer cells often exhibit large-scale genomic rearrangements, including copy number variation (CNV) as well as structural variations (SVs), which have been implicated in the initiation and progression of cancer^41^. In previous studies, Hi-C has been shown to detect these genomic variations in bulk tumor tissues with high accuracy and efficiency^42, 43^. However, these bulk analyses could not resolve the heterogeneity and evolution of tumor populations in response to therapies. Droplet Hi-C overcomes this problem through detection of genomic variations in individual cancer cells. As a proof of principle, we performed Droplet Hi-C with two colorectal cancer cell lines, COLO320DM and COLO320HSR, derived from the same patient, both carrying similar copies of *MYC* amplicon. In COLO320DM, the *MYC* oncogene is located on ecDNA, while In COLO320HSR, it exists on chromosomes as tandem repeats, termed homogeneously staining regions (HSR). By inferring the copy number of each 1-Mb bin in single cells as well as aggregated pseudo-bulk profiles, we detected elevated copies of the *MYC* locus from both samples, as anticipated (Fig. 3a). We also observed fluctuations in the copy numbers across the entire genome (Fig. 3a). We used the inferred DNA copy number of each genomic region to correct bias in the contact matrices and then identified SVs. Our analysis revealed distinct SVs between these two cancer lines, and we showed one example of duplication event on chromosome 6 (Fig. 3b).

**Fig. 3:**
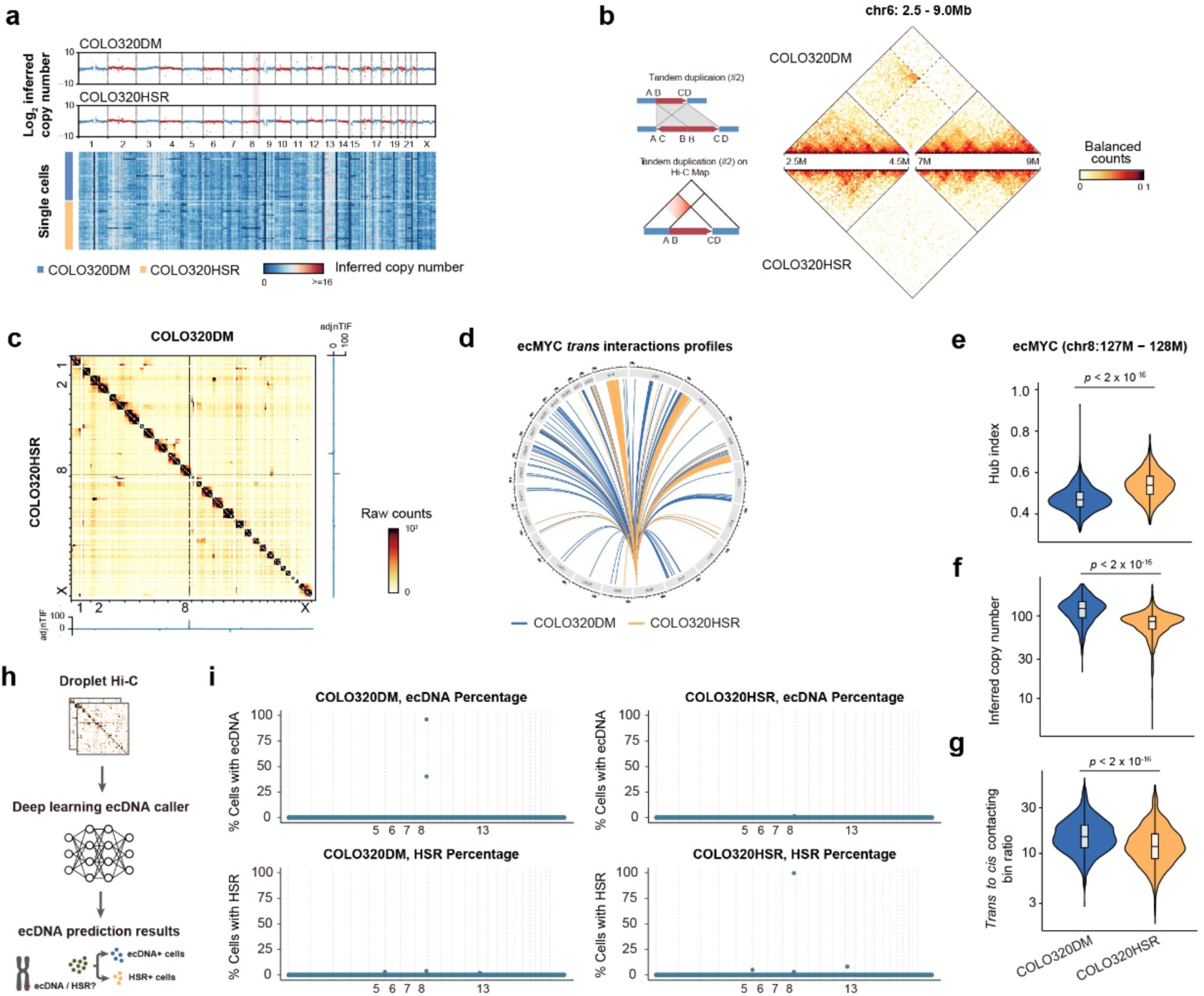
Droplet Hi-C illuminates copy number, structural variations and ecDNA in tumor samples. **a,** Copy number inferred from pseudo-bulk Droplet Hi-C profiles in COLO320DM and COLO320HSR, with heatmap showing representative single-cell CNV in each sample. The ecMYC bins in COLO20DM are highlighted in pink. **b,** Example of a sample-specific SV on chromosome 6, predicted with EagleC. Cartoon illustration explaining the rearranged contacts pattern is shown aside. **c,** Comparison of genome-wide contact maps and adjusted normalized inter-chromosomal interaction frequency (adj_n_TIF) between COLO320DM and COLO320HSR. **d,** Circos plot showing *trans*-interaction profiles of genomic bin containing *MYC* in COLO320DM or COLO320HSR. **e-g,** Distribution of single-cell hub index **(e)**, copy number **(f)**, and *trans*-to-*cis* contacting bin ratio **(g)** of ecMYC in COLO320DM and COLO320HSR. n = 1,352 (Hub index, COLO320DM), 1,366 (Hub index, COLO320HSR), 1,426 (Inferred copy number and *trans*-to-*cis* contacting bin ratio, COLO320DM), 1,535 (Inferred copy number and *trans* to *cis* contacting bin ratio, COLO320HSR); *P* values are from Wilcoxon signed-rank test. **h,** Schematics of deep learning-based ecDNA caller. **i,** Dot plot showing genome-wide ecDNA and HSR prediction results from deep learning-based ecDNA caller for COLO320DM and COLO320HSR cells.

It has been difficult to distinguish whether the *MYC* exists on the ecDNA or as HSR from whole genome DNA sequencing alone. We ran AmpliconArchitect algorithm using whole-genome sequencing (WGS) data of COLO320HSR, and AmpliconArchitect classified the HSR as a circular amplicon. While the genome-wide interaction patterns have been proposed as potential indicators of ecDNA^44^, aggregate Hi-C profiles showed similar patterns of inter-chromosomal interactions between the amplified *MYC* locus and the rest of the genome in both COLO320DM and COLO320HSR (Fig. 3c). Although measurement like adjusted normalized inter-chromosomal interaction frequency (adj_n_TIF) has been proposed to determine ecDNA identity^44^, we found that adj_n_TIF was equally high at the 1-Mb bin containing *MYC* in both cell lines, indicating that this value is likely driven by abnormally high copy number rather than other specific property of ecDNA (Fig. 3c).

We therefore tried to discern distinctions between *MYC* ecDNA and HSR by analyzing their interactome obtained from the Droplet Hi-C dataset. In COLO320DM, relatively uniform interactions happened within ecDNA (Extended Data Fig. 4a). In COLO320HSR, we found two deletions, evidenced by significantly lower numbers of sequencing reads at those regions. Single-cell profiles confirmed that these two regions appeared uniformly across all the cells (Extended Data Fig. 4a). Interestingly, we found that ecDNA interacts with other chromosomes more evenly, while HSR displayed a preference towards certain chromosomes (Fig. 3d). To quantitatively measure the evenness of inter-chromosomal contacts, we devised the hub index as the Gini coefficient of inter-chromosomal contacts for the 1 Mb *MYC* bin in each single cell. COLO320DM shows a significantly smaller hub index, indicating a more uniform *trans*-interacting profile (Fig. 3e). We also quantified other parameters which can be obtained from single-cell Hi-C data, including inferred copy number (Fig. 3f) and the *trans*-to-*cis* contacting bin ratio (Fig. 3g. Both of these parameters exhibited significant differences between COLO320DM and COLO320HSR (Fig. 3f-g). COLO320DM cells had approximately 47% more copies than COLO320HSR cells at the *MYC* genomic bin, which aligns with previous research findings^45^ (Fig. 3f). Additionally, COLO320DM displayed a higher *trans*-to-*cis* contacting ratio for the *MYC* bin compared to COLO320HSR (Fig. 3g). These results collectively suggest that *MYC* ecDNA exhibits distinct interaction patterns from *MYC* HSR at the single-cell resolution, making it a potential indicator for determining whether a genomic bin is likely to be ecDNA or not within an individual cell.

The measurement of *trans*-interaction evenness also paves the way to investigate whether multiple ecDNA tend to cluster into “hubs” in a single cell, since the agglomerated ecDNAs will show contacting preference against specific chromosomes compared to diffused ecDNAs. By shuffling the inter-chromosomal contacts across genome to generate an uniform distribution as reference for the 1Mb *MYC* bin in COLO320DM and COLO320HSR, we found that the hub index of inter-chromosomal contacts in COLO320HSR is significantly higher, demonstrating that this genomic bin showed stronger bias in contacts for HSR, consistent with the tandem repeat aggregation property of HSR (Extended Data Fig. 4b). However, COLO320DM exhibits a slightly lower hub index compared to the random background, indicating that ecDNA does not have a strong nuclear localization bias in these cells (Extended Data Fig. 4b), and that previously reported ecDNA hubs in COLO320DM are likely transient or infrequent features^46^.

The observed differences in contact profiles between ecDNA and HSR hint at potential ways to distinguish these aberrant chromosomal features from single-cell Hi-C data. To this end, we fit a multivariate logistic regression model using hub index, estimated copy number, *trans*-to-*cis* contacting bin ratio for ecDNA identification. This regression model is then used to predict ecDNA probability for each 1Mb bin that contains ecDNA among the entire cell population, as well as whether each 1Mb bin contains ecDNA in each single cell (Extended Data Fig. 5a). Specifically, the *MYC* locus (chr8:127Mb-128Mb) in COLO320DM cells is treated as ground truth and in COLO320HSR cells as negative in the training dataset. On the validation dataset, the model reached a 0.99 specificity, a 0.97 precision, a 0.89 accuracy and a 0.71 recall. We first applied the regression model to 1,426 COLO320DM cells and 1,535 COLO320HSR cells, which identified two consecutive bins (chr8:126Mb-128Mb) as ecDNA. In COLO320DM, 72.7% of cells were predicted to be ecDNA positive (Extended Data Fig. 5b). In contrast, only 20.5% of cells were predicted to be ecDNA positive in COLO320HSR cells (Extended Data Fig. 5b). We also analyzed the 3,423 cells obtained from the adult mouse cortex, and found that no bin contains ecDNA. Taken together, these results show that our logistic regression model based ecDNA caller algorithm can identify known ecDNA from COLO320DM, although the false positive rate remains a problem in the COLO320HSR samples.

To further improve the accuracy of our ecDNA caller, we developed a deep learning model based on convolutional neural networks (CNN) (Fig. 3h). The deep learning model utilizes the sliced contact matrix, comprising five consecutive 1Mb genomic bins as its input. At the heart of this process, a neural network is trained using these input variables to identify ecDNA within each central bin, providing predictions for the presence of ecDNA or HSR in 1Mb bins, both at the cell population and single-cell levels. Our deep learning model has demonstrated remarkable performance, achieving a recall rate of 0.80, while maintaining a 0.99 precision in ecDNA prediction, surpassing that of the logistic regression model. Furthermore, it achieves 0.99 specificity and 0.93 accuracy, as well as a faster processing speed. By applying the deep learning-based ecDNA caller to the COLO320DM and COLO320HSR validation dataset, we were able to identify *MYC* ecDNA in 96% of COLO320DM cells and HSR in 3.86% of COLO320DM cells (Fig. 3i, Supplementary Table 3). Conversely, MYC HSR was identified in 0.20% of COLO320HSR cells, along with HSR in 99.54% of COLO320HSR cells (Fig. 3i). Given that the deep learning-based ecDNA caller pipeline can directly distinguish between ecDNA and HSR, while also outperforming the regression model-based algorithm, we decided to employ the deep learning model-based ecDNA caller algorithm in all the subsequent studies.

### Droplet Hi-C uncovers ecDNA heterogeneity and tumor cell evolution in response to drug treatment

Previous studies showed that glioblastoma (GBM) cells carrying extrachromosomal oncogenic variant of epidermal growth factor receptor (EGFRvIII) can develop drug resistance after treatment with tyrosine kinase inhibitors such as erlotinib (ER), which coincides with the loss of *EGFR* ecDNA^47^. To further characterize the dynamics of *EGFR* ecDNA abundance and heterogeneity during the acquisition of drug resistance, we employed Droplet Hi-C to profile both naive GBM39 cells and cells after more than 30-days of treatment of erlotinib (GBM39-ER) (Fig. 4a). With 9,204 high-quality nuclei profiled, we were able to resolve five different clusters of cells based on chromatin architecture in each cell (Fig. 4b, Extended Data Fig. 6a). After erlotinib treatment, a notable evolutionary shift is observed in the GBM39 cells. Subpopulation cluster 2 (C2) nearly vanished, subpopulation cluster 0 (C0) expanded, and several new subpopulations including clusters 1, 3, and 4 (C1, C3, C4) emerged (Fig. 4c-d).

**Fig. 4:**
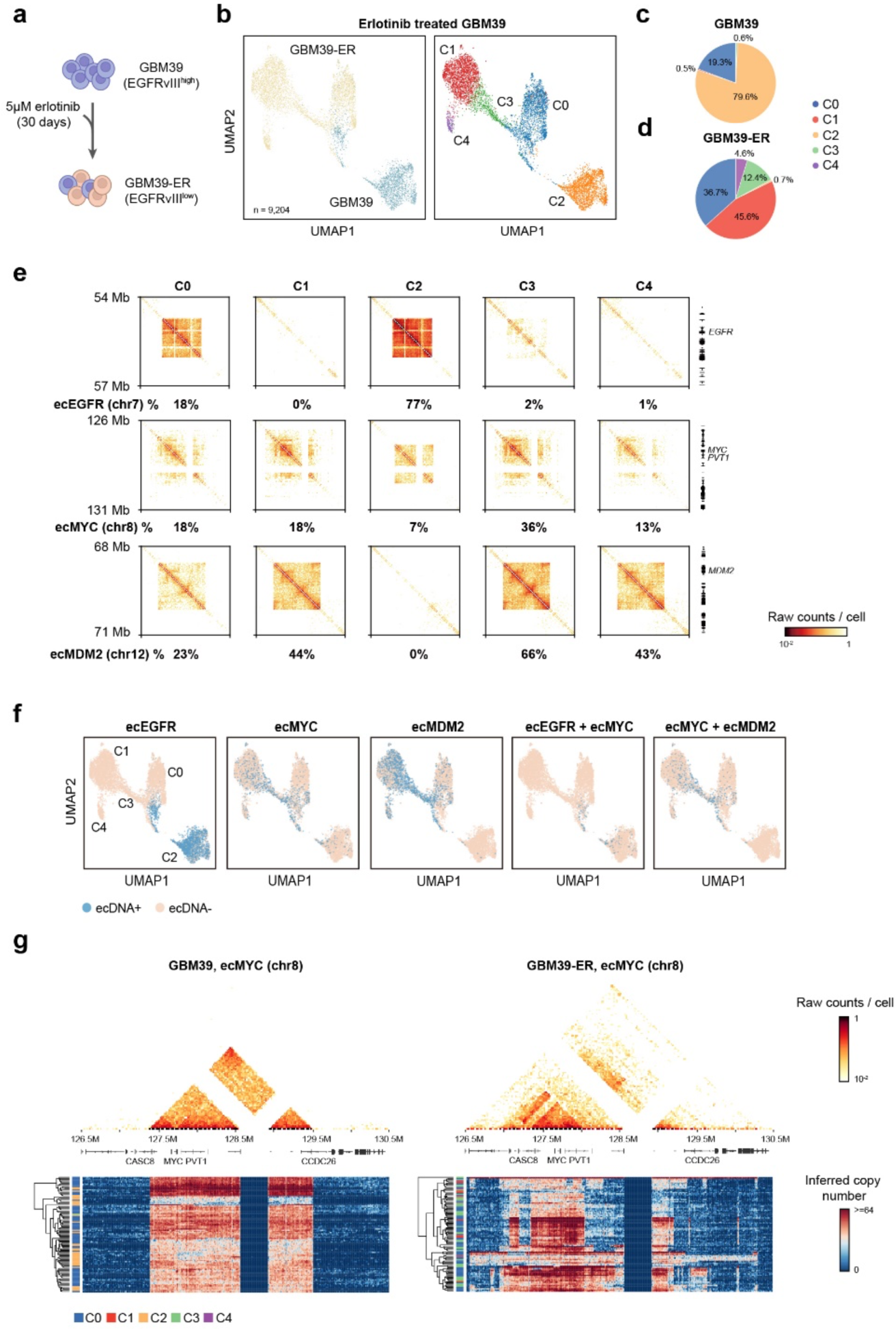
Droplet Hi-C reveals ecDNA dynamics under drug treatment. **a,** Illustration of the erlotinib treatment procedure for GBM39 cells. **b,** UMAP embedding visualization and clustering analysis of Droplet Hi-C data from GBM39 before- and after-erlotinib treatment. **c-d,** Pie charts showing the cell proportion of each cluster in GBM39 **(c)** and GBM39-ER **(d)**. **e,** Comparison of contact maps and percentages of ecDNA-positive cells among all clusters for ecEGFR, ecMYC and ecMDM2. Genomic coordinates for key oncogenes are displayed aside. **f,** UMAP visualization of the ecDNA positive cell distribution within each cluster. **g,** Pseudo-bulk contacts map at 10-kb resolution showing ecMYC local structure in both GBM39 and GBM39-ER samples. Heatmaps representing representative single-cell inferred CNV in the same genomic range are shown below.

At the sample level, elevated inter-chromosomal contacts are observed for *EGFR* locus and, surprisingly, *MYC* locus before treatment. After erlotinib treatment, the extensive inter-chromosomal contacts at *EGFR* locus diminished, but increased at the *MYC* locus and a new locus on chromosome 12 encompassing *MDM2* (Extended Data Fig. 6b). We utilized our ecDNA caller to identify candidate ecDNAs in the Droplet Hi-C data from both GBM39 and GBM39-ER cells (Supplementary Table 3). In GBM39, we could detect ecDNA containing the *EGFR* locus (ecEGFR) and ecDNA containing the *MYC* locus (ecMYC) (Extended Data Fig. 6c). We also detected a low-frequency ecDNA on chromosome 18 (1.5%, ecChr18) (Extended Data Fig. 6d). In GBM39-ER cells, we discovered ecDNA harboring the *MDM2* locus (ecMDM2) and also ecMYC (Extended Data Fig. 6c).

At the cluster level, we showed the percentage of cells containing these three ecDNA candidates in every cluster, revealing dynamics of ecDNA-harboring cancer cell populations during development of drug resistance (Fig. 4e). Specifically, ecEGFR disappeared nearly completely after erlotinib treatment (C1, C3, C4) as previously reported^47^, whereas ecMDM2 appeared only in erlotinib-resistant populations and is dominant in GBM39-ER subgroup (C3). And ecMYC showed low frequency in GBM39 population (C2) and increased in transition state clusters (C0, C1, C3). We also plotted the contact maps at 25-kb resolution for these three ecDNAs in every cluster. We observed that the structure and size of the ecMYC amplicon were altered, unlike ecEGFR and ecMDM2, suggesting a more intricate transformation at the *MYC* ecDNA during the drug treatment (Fig. 4e).

Surprisingly, when we plotted the distribution of each ecDNA among all the identified clusters (Fig. 4f), we found coexistence of distinct ecDNA species in some tumor cells (Extended Data Fig. 6e). In cluster C0, a small population of cells (4.7%) harbored both ecMYC and ecEGFR. These cells may potentially serve as the drug-resistant pool of cells during erlotinib treatment, given that the gain of ecMYC may increase cellular fitness. This observation raises the intriguing question of whether GBM39-ER was selected from a small population of cells by erlotinib treatment. Since the simultaneous increase in copy number can indicate whether two adjacent genomic regions are co-present in the same ecDNA, we focused on the ecMYC and characterized its structure in GBM39 cells before and after erlotinib treatment at 10-kb resolution. Although the copy number varies across different cells, the boundaries of ecMYC in GBM39 are almost identical (Fig. 4g). Strikingly after treatment, the ecMYC boundaries as well as internal structure displayed huge variability, reflecting the heterogeneity of MYC ecDNA at single-cell resolution. Boundary variations in ecDNA did not exhibit specificity with relation to clusters (Fig. 4g). We next selected the same number of cells from either GBM39 or GBM39-ER within the transition cluster (C0) and compared the aggregated contacts map at ecMYC regions to see whether an intermediate state for ecMYC structure can be found. Both profiles, although from the same cluster, still resembled their original samples and showed the same differences in ecMYC structure, instead of showing a similar intermediate structure (Extended Data Fig. 6f). Therefore, we reasoned that for GBM39 cells, erlotinib treatment induced the formation of unique cell populations containing distinct *MYC* ecDNA, instead of selectively enriching a pre-existing cell population harboring ecMYC with growth advantage upon treatment.

### Droplet Hi-C detects ecDNA in primary tumor samples

ecDNA is not only a biomarker for poor prognosis but also a crucial driver in GBM^48^. As Droplet Hi-C successfully revealed dynamic changes of ecDNA in GBM39 cells associated with drug treatment, we further sought to apply Droplet Hi-C to an IDH-wildtype GBM tumor sample (Fig. 5a). By employing only chromatin structure information for cell embedding, we segregated the cells into tumor-like and non-tumor cells (Fig. 5b), featuring expected CNV events in tumor-like cells such as loss of chromosome 10 and extensive amplification of chromosome 7 (Fig. 5c). We found that the genomic region with the most variable copy numbers on chromosome 7 carried the oncogene *EGFR* (Fig. 5c-d), and also displayed an ecDNA-like contact pattern in the contact map of the tumor-like population (Fig. 5e). Utilizing our ecDNA caller, the tumor-like population had significant enrichment of ecEGFR as expected (Fig. 5f, Supplementary Table 3).

**Fig. 5:**
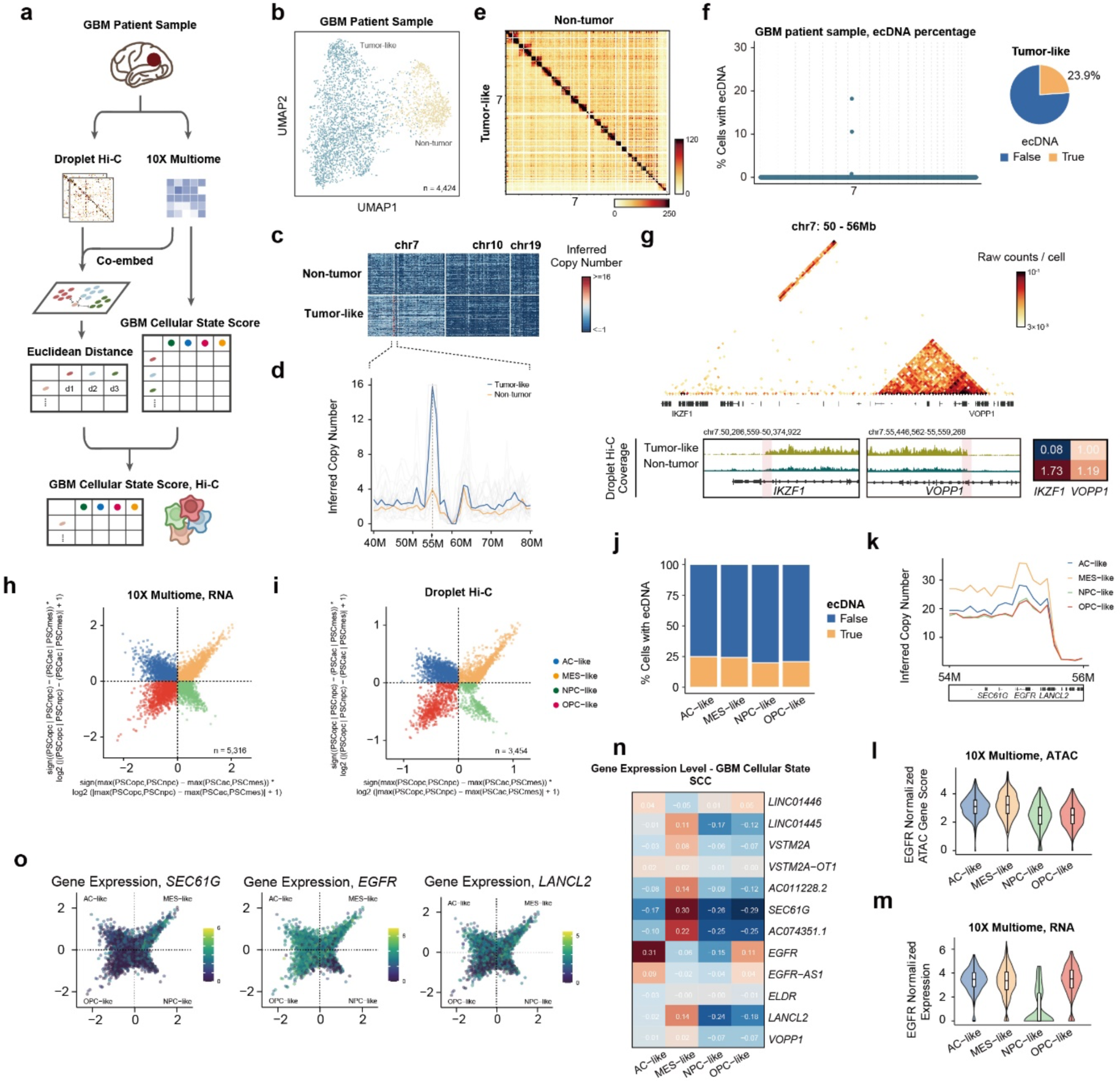
Droplet Hi-C depicts ecDNA variations in patient samples. **a,** Schematics of GBM cellular state analysis for Droplet Hi-C data by co-embedding with reference 10x Multiome data. **b,** UMAP visualization of Droplet Hi-C data on GBM patient sample. **c,** Representative single-cell inferred CNV on chromosome 7, 10 and 19 in tumor-like and non-tumor populations identified from Droplet Hi-C. **d,** Line plot showing copy number among single cells in tumor-like and non-tumor populations. Lines with color highlighted the median profiles. The single-cell copy number profile examples are colored in gray (n = 20). **e,** Genome-wide contact maps from tumor-like and non-tumor populations. Color bar shows the raw contact number. **f,** The ecDNA prediction results in GBM patient sample. The percentage of cells predicted to contain ecEGFR in tumor-like population is shown in a pie chart. **g,** An example of tumor-like population-specific SV. Genome browser view showing the Droplet Hi-C read coverage at associated genes displaying below. Predicted breakpoints of SV are highlighted in pink. Gene expression level (RPKM) of associated genes in different populations from 10x Multiome is also shown. **h-i,** Two-dimensional representation of cellular states based on scRNA-seq data from 10x Multiome **(h)** and Droplet Hi-C data **(i)**. Each quadrant corresponds to one cellular state. **j,** The percentage of ecEGFR-positive cells across four different GBM cellular states. n = 563 (AC-like), 600 (MES-like), 247 (NPC-like), 346 (OPC-like). **k,** Inferred copy number at 100-kb resolution of regions on ecEGFR among different GBM cellular states. **l-m,** Violin plots showing *EGFR* ATAC gene score **(l)** and *EGFR* gene expression level **(m)** across four different GBM cellular states from 10X Multiome dataset. Only cells with GBM cellular states score pass cutoff (± 0.5) are shown. n = 244 (AC-like), 410 (MES-like), 125 (NPC-like), 263 (OPC-like). **n,** Heatmap showing Spearman’s correlation coefficients (SCC) between GBM cellular states score and gene expression level for all the genes located on ecEGFR. **o,** Two-dimensional representation of GBM cellular states, colored by ecDNA genes expression levels.

In addition to CNV and ecDNA, Droplet Hi-C also identified tumor-like cell-specific chromatin organization and SVs in the patient sample (Fig. 5g, Extended Data Fig. 7a-b). Based on the sequencing reads coverage and abnormally high interaction frequency, *IKZF1* gene, encoding a tumor suppressor, was found to translocate to ecEGFR without the TSS sequence, leading to its transcriptional disruption in the tumor cell population (Fig. 5g). Among the genes located on the ecDNA, the *VOPP1* gene exhibited the strongest interaction frequency with *IKZF1*, suggesting their fusion (Fig. 5g). Interestingly, this fusion event did not show any discernible effect on the transcriptional activity of *VOPP1*, based on the aggregated expression profiles from single-cell RNA-seq data (Fig. 5g).

Previous single-cell RNA-seq profiling of gliomas defined four major cellular states for tumor cells, including neural progenitor-like (NPC-like) cells, oligodendrocyte progenitor-like (OPC-like) cells, astrocyte-like (AC-like) cells and mesenchymal-like (MES-like) cells^49^. To investigate whether ecDNA is associated with specific cellular states, we obtained single-cell gene expression together with chromatin accessibility at single-cell resolution using 10X multiome. We co-embedded the Droplet Hi-C data with single-cell RNA-seq, and used k-nearest neighbors to assign one of the four malignant cellular states to individual cells from the Droplet Hi-C dataset (Fig. 5h-i, Extended Data Fig. 7c). We subsequently characterized the differential ecDNA features across these states. We found that the fractions of cells containing ecEGFR were similar among the four cell states (Fig. 5j), but the cells in AC- and MES-like states contained higher copy numbers of ecEGFR (Fig. 5k). The increase of copy number was associated with higher *EGFR* expression level and chromatin accessibility in these states (Fig. 5l-m). Furthermore, expression levels of genes on the ecDNA showed strong correlation with the cellular states score (Fig. 5n). Consistent with previous reports^49^, the expression level of *EGFR* exhibited a high correlation with AC-like state scores (Fig. 5n). We also found that genes on the same ecDNA displayed varying correlation levels with different cell states, and that such correlation could not be fully explained by ecDNA copy number, indicating a complex regulatory mechanism governing ecDNA genes expression across cellular states. Examples besides *EGFR* included the preferential expression of known *EGFR* co-amplified gene *SEC61G* in MES-like state, and the preferential expression of *LANCL2* in AC- and MES-like states (Fig. 5o). The abundance of ecDNA seems to be more closely related to the differentiation states (AC- and MES-like), although mechanism dictating varied expression levels of ecDNA genes besides copy number effects requires further investigation.

To further demonstrate the utility of Droplet Hi-C in clinical tumor samples, we adopted it to analyze the bone marrow mononuclear cells (BMMC) from an acute myeloid leukemia (AML) and myelodysplastic syndrome (MDS) patient before and after treatment with azacitidine and ventoclax. Azacitidine is a DNA methyltransferase inhibitor known to slow the growth of AML cells in part by promoting their differentiation into more mature cells (Extended Data Fig. 7d). Venetoclax is an inhibitor of the anti-apoptotic protein BCL2. The patient entered a remission from secondary AML after completing one cycle of therapy. The contact maps revealed that the before-treated BMMC initially harbored a strikingly large (∼5 MB) ecDNA containing the *MYC* gene. After treatment, the ecMYC can no longer be detected by Droplet Hi-C (Extended Data Fig. 7e). To classify tumor-like and non-tumor cells in BMMC, we leveraged known mutations detected by targeted bulk DNA sequencing to distinguish malignant cells. After drug treatment, the proportion of cells harboring mutations also diminished (Extended Data Fig. 7f). Clinical karyotype analysis showed the persistence of trisomy 8 which was present while the patient had MDS prior to AML transformation. Thus, we speculated that remaining mutant reads might originate from pre-leukemic MDS cells.

Expression of the oncogene *MYC* is up-regulated by numerous tissue-specific enhancers through long-range interactions in different cancers^50^. One of the evolutionarily conserved enhancer clusters (blood enhancer cluster, BENC), which is ∼1.8 Mb downstream of *MYC*, has been shown to activate *MYC* expression in hematopoietic stem cells and AML^50^. The long-range interactions between *MYC* and BENC were observed in the before-treated BMMC sample. The detection of focal amplification of BENC is also consistent with a previous report^50^ (Extended Data Fig. 7g). Specifically, both the *MYC* gene and BENC were located on the ecMYC. The interaction between *MYC* and BENC is obvious in the before-treated sample and disappears after treatment, suggesting its role in tumor initiation and progression (Extended Data Fig. 7h). Thus, in this patient’s tumor, ecDNA may promote *MYC* expression not only through increased DNA copy number but also cancer-specific enhancer-promoter interactions.

In summary, Droplet Hi-C facilitates the detection of ecDNA dynamics in tumors before and after drug treatment. Besides ecDNA, identification of canonical chromatin structures such as SVs and chromatin loops also helps to elucidate the ectopic regulatory programs responsible for tumor progression and drug resistance. Droplet Hi-C therefore shows great potential for advancing understanding of mechanisms driving tumor clonal evolution and sensitivity to treatment.

### Droplet-based joint profiling of chromatin architecture and transcriptome

Single-cell joint profiling of chromatin organization and transcriptome can facilitate the investigation of the relationships between gene expression and chromatin architecture. We modified the above Droplet Hi-C protocol to be compatible with the 10x Genomics Chromium Single Cell Multiome kit, to achieve simultaneous profiling of RNA and Hi-C from single nuclei (Fig. 6a, Extended Data Fig. 8), termed Paired Hi-C. Like Droplet Hi-C, Paired Hi-C also offers benefits in terms of throughput and hands-on time over other microwell- or combinatorial indexing-based joint Hi-C/RNA-seq single-cell assays (Extended Data Fig. 8).

**Fig. 6:**
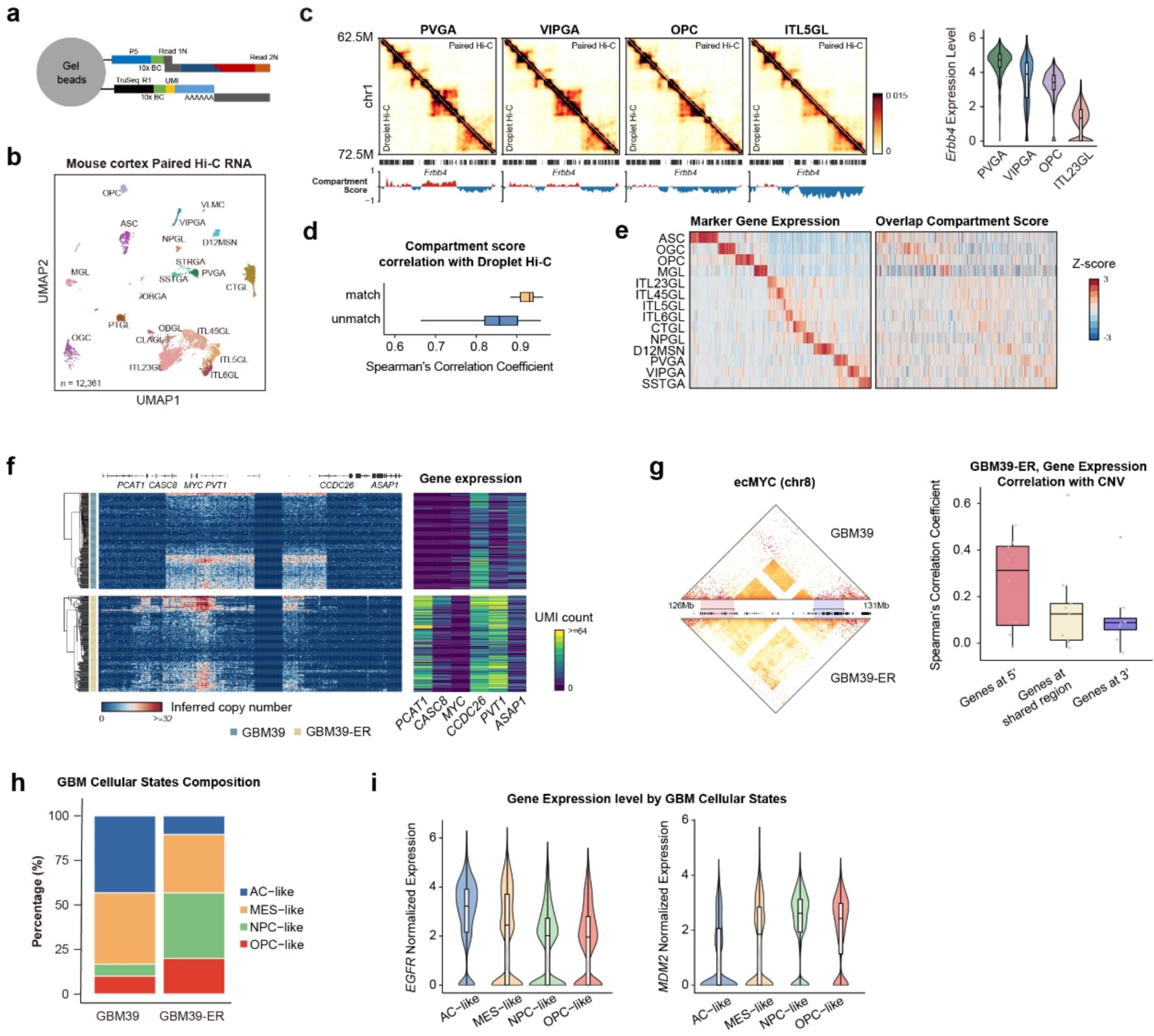
Joint profiling of single-cell Hi-C and transcriptome. **a,** Schematic of the Paired Hi-C molecule barcoding step with 10X Multiome kit. **b,** UMAP visualization of mouse cortex single-cell transcriptome from Paired Hi-C. **c,** Comparison of pseudo-bulk contact maps between Droplet Hi-C and Paired Hi-C at the region surrounding gene *Erbb4,* along with compartment score profiles from Paired Hi-C in representative cell types. Color bar shows the imputed contact number. A violin plot of *Erbb4* expression level in representative cell types is also shown. **d,** Comparison of Spearman’s correlation coefficients (SCC) between compartment score across cell types in Droplet Hi-C and Paired Hi-C. n =15 (match); 225 (unmatch). **e,** Heatmap showing marker genes expression levels and compartment scores of corresponding bins among all clusters. **f,** Single-cell inferred copy number heatmaps of regions harboring ecMYC in GBM39 and GBM39-ER. Single-cell UMI counts heatmaps of representative genes on ecDNA are also shown. **g,** SCC between gene expression and copy number for ecMYC genes in 5’, 3 ‘variable regions and shared regions in GBM39-ER. Illustration of ecMYC variable regions is shown on the left. n = 9 (5’ gene); 10 (shared); 11 (3’ gene). **h,** Comparison of cell proportions in four GBM cellular states between GBM39 and GBM39-ER samples. **i,** Comparison of expression levels of *EGFR* and *MDM2* among four GBM cellular states. n = 4,307 (AC-like), 6,791 (MES-like), 4,945 (NPC-like), 3,123 (OPC-like).

To demonstrate the utility of Paired Hi-C, we applied it to a cortex specimen collected from an 8-week old mouse. Overall, we obtained 12,361 joint single-cell profiles of transcriptome and chromatin structure with 42,210 chromatin contacts per cell (duplication rate 66.2%), and a median of 3,914 UMI per cell and 1,746 genes per cell (duplication rate 74%, Extended Data Fig. 9a). From Paired Hi-C RNA modality, we successfully identified 20 major cell types in the mouse cortex, including 9 excitatory neuron subtypes, 6 inhibitory neuron subtypes, and 5 non-neuronal subtypes (Fig. 6b, Supplementary Table 4). The annotation is transferred from Droplet Paired-Tag. The accuracy of cell type identification was confirmed by overlapping with other single-nuclei RNA-seq datasets and the unique expression patterns of marker genes (Extended Data Fig. 9b-d). While single-cell 3D genome features were relatively limited in Paired Hi-C, the compartment score computed from Paired Hi-C data for each cell type (as defined by single-nuclei RNA-seq) exhibited a stronger correlation with the matched cell type’s Droplet Hi-C and sn-m3C-seq data (Fig. 6c-d, Extended Data Fig. 9e). At cell type-specific marker genes, we can also detect concordant changes of compartment and gene expression (Fig. 6e). As an example, *Erbb4* genes were located within the A compartment in PVGA and VIPGA GABAergic neurons, while they were placed in the B compartment in OPC non-neuronal types and ITL5GL glutamatergic neurons. The expression level of *Erbb4* was notably higher in PVGA and VIPGA than OPC and ITL5GL (Fig. 6c). We also utilized transcriptome information from Paired Hi-C and co-embed it with Droplet Hi-C. The results agree well with the cell annotations based on Droplet Paired-Tag (Extended Data Fig. 9f). As such, we can augment the number of chromatin contacts for each cell type by combining Paired Hi-C and Droplet Hi-C.

The heterogeneity in the ecMYC boundaries at single-cell level within GBM39-ER prompted the question of whether this variable ecMYC boundary could result in varying ecDNA gene expression within individual cells. By applying Paired Hi-C in GBM39 and GBM39-ER cells, we analyzed ecDNA structure and the expression level of its related genes in single cells. We found that in general, the genes located on the ecMYC regions within GBM39 and GBM39-ER displayed elevated expression levels where their underlying copy number surges, indicating a higher probability of being part of ecMYC (Fig. 6f). For genes located 5’ ecMYC boundaries such as *CASC8* and *PCAT1*, their expression exhibited a strong correlation with the copy number of underlying DNA segments (Fig. 6f). In the case of ecMYC constant genes, such as *PVT1*, expression was constantly high and also positively correlated with local contact numbers (Fig. 6f). Unexpectedly, the expression of the *CCDC26* gene, located at the 3’ end of the ecMYC variable region, was high in GBM39-ER cells despite low copy number (Fig. 6f). When we conducted correlation analysis between gene expression levels and copy numbers for the genes at the 5’ and 3’ ends of the variable ecMYC boundary separately in GBM-ER cells, the findings revealed that genes located within the 5’ ecMYC variable region exhibited a more pronounced increase in expression with rising copy numbers. In contrast, genes within the 3’ ecMYC variable region showed much lower correlations between gene expression levels and copy numbers (Fig. 6g). These results highlight the complex relationship between amplicon structure and gene expression.

Besides ecMYC structural heterogeneity and gene expression variation, we found that ecMYC displayed different *trans*-interaction patterns in GBM39 and GBM39-ER cells. These different *trans*-interaction patterns of ecDNA were also associated with distinct transcriptional responses. Specifically, we observed that GBM39-specific interacting genes showed higher expression levels in GBM39 than in GBM39-ER cells, while GBM39-ER-specific interacting genes had higher expression levels in GBM39-ER cells (Extended Data Fig. 10a). This suggests a potential mobile regulatory role of ecDNA in global chromosomal transcription^44^.

In the GBM patient sample, we found that ecEGFR showed higher copy numbers in AC-like and MES-like cell populations as well as higher expression levels of *EGFR*, which may render the cellular states changes under erlotinib treatment. Using previously defined methods, we utilized Paired Hi-C single-cell RNA-seq data to classify GBM39 and GBM39-ER cells into four GBM cellular states, and found a significant reduction of differentiated-like population after erlotinib treatment (Fig. 6h). We previously demonstrated that GBM39-ER cells were characterized by the loss of ecEGFR and the emergence of ecMDM2, and we sought to determine whether the expressions of these genes enriched in particular states within the resistant cell population. As found in the GBM tumor sample, *EGFR* expression was higher in AC-like and MES-like cells (Fig. 6i). Interestingly, *MDM2* expression was enriched in the progenitor states (OPC- and NPC-like), which became more prevalent after treatment (Fig. 6i). These results highlight the unique interplay between ecDNA dynamics, transcriptionally defined cellular states, and drug resistance.

Besides ecDNA, chromatin architecture also undergoes a dramatic shift during erlotinib treatment, along with changes in gene expression. In GBM39 cells following erlotinib treatment, we observed 1,066 A compartments transitioned into B compartments, whereas 2,796 B compartments shifted to A compartments. The majority of compartment identities remained stable throughout the treatment, including 10,930 A compartments and 11,876 B compartments (Extended Data Fig. 10b). This compartment shift is also accompanied with gene expression changes. Specifically, genes situated within compartments that changed from A to B exhibited a significant decrease in expression, while those transitioning from B to A showed a significant increase in expression. In contrast, gene expression within stable compartments displayed only slight expression alterations (Extended Data Fig. 10c).

In summary, by incorporating transcriptomic data with chromatin structure information, Paired Hi-C enables direct interrogation of the effects of chromatin reorganization and structural alterations like ecDNA on gene expression, providing a valuable tool for characterizing gene regulatory programs in development and disease pathogenesis.

## Discussion

In recent years, there has been a significant surge in the development and application of single-cell Hi-C methods. However, current single-cell Hi-C methods still face challenges when applying to heterogeneous tissues and tumor biopsies. Our Droplet Hi-C leverages the commercially available microfluidic platform to provide high-throughput and fast single-cell Hi-C assays with minimal hands-on time, and relatively low costs. Utilizing Droplet Hi-C technology on the adult mouse cortex, we identified cell type-specific hierarchical chromatin structures and established correlations with epigenetic modifications. Additionally, our study unveiled alterations in chromatin organization, CNV, SVs, and ecDNA in cancer cells and tumor patient samples, especially during drug treatment. We further extend Droplet Hi-C to Paired Hi-C, which achieves joint profiling of chromatin architecture and transcriptome in single cells. This enables the association of gene expression phenotypes with 3D genomic structure. The complexity of the Hi-C modality in Paired Hi-C is notably lower when compared to Droplet Hi-C. Nonetheless, we have shown that it is possible to leverage transcriptome data obtained from Paired Hi-C experiments for co-embedding with Droplet Hi-C data. This approach enabled us to resolve and annotate cell types and supply Hi-C contact information for each cell type to enhance the resolution of chromatin organization.

EcDNA was first identified in 1965, described as double minutes (DM), and is prevalent in cancer as they often harboring oncogenes^51, 52^. Unlike the kilobase-sized circular DNA (eccDNA) found in healthy somatic tissues^53^, ecDNA varies in size from dozens of kilobases to megabases, making it 100 to 1,000 times larger^54^. The ecDNA is characterized by high amplification and a circular structure^55^. Current single-cell ecDNA detection methods can be broadly categorized into two primary groups. The first group involves the use of computational tools and WGS data obtained from NGS^56^ or TGS^57, 58^ to identify circularization sites. However, distinguishing between ecDNA and HSR using this approach can be challenging, because they have the same breakpoint sequences. The second group is based on Circle-seq^59–61^, a strategy that utilizes exonuclease to degrade linear DNA to retain circular DNA for sequencing. This method allows for the differentiation of ecDNA from linear tandem repeats. However, Circle-seq-based strategies have been associated with a high false-positive rate due to incomplete degradation of linear DNA and multiple displacement amplification (MDA). Accurate identification of ecDNA using its multiple features at single-cell level remains challenging. Here, we have developed an innovative ecDNA detection algorithm that utilizes both *trans* and *cis* interactions data from Droplet Hi-C datasets, enabling precise ecDNA identification while effectively distinguishing it from HSR. With our deep learning-based ecDNA caller algorithm, we can reliably detect ecDNA in both cancer cell lines and clinical tumor samples in each single cell, and calculate their proportion within the cell population. Furthermore, the increased occurrence of intra-ecDNA contacts provides valuable support for pinpointing ecDNA boundaries at the single-cell level. For instance, we observed variations in the ecMYC boundary within the GBM39-ER cell population. Additionally, we are capable of detecting SVs events within both ecDNA and HSR.

Pan-cancer studies have demonstrated the detection of ecDNA in numerous human cancer types and that ecDNA is associated with inferior clinical outcomes across all amplification classes^62–66^. Further, new studies have begun to highlight unique therapeutic vulnerabilities of tumors that harbor ecDNA^67, 68^. Thus, given the independent prognostic value of ecDNA and its potential to be targeted, recent calls have been made to include ecDNA profiling in clinical specimens^69^. Droplet Hi-C and deep learning-based ecDNA caller algorithm offer a reliable toolkit for the identification of ecDNA in clinical samples as demonstrated in patient GBM and AML tumors. This toolkit may be particularly useful in heterogeneous patient samples that harbor ecDNA species whose structures and copy numbers may evolve subclonally through treatment. Importantly, our ecDNA caller algorithm possesses the unique capability to differentiate between amplifications occurring on ecDNA and HSRs. Such differentiation may carry critical therapeutic and clinical implications. In our study, we observed the disappearance of ecEGFR, emergence of ecMDM2 and structural dynamics of ecMYC in GBM39 cells following treatment with erlotinib, a tyrosine kinase inhibitor commonly used in the management of *EGFR*-positive tumors. Notably, the effectiveness of erlotinib treatment in GBM was poor, possibly attributed to the ecDNA evolution. Additionally, in an AML patient sample, we noted that ecMYC could no longer be detected after treatment with azacitidine and venetoclax. Collectively, our methods can be used in ecDNA detection in clinical samples and hold promise for advancing research related to ecDNA in tumor diagnosis and treatment.

## Methods

### Cell culture

HeLa S3, K562, GM12878, COLO320DM, and COLO320HSR cells were cultured in high-glucose DMEM or RPMI-1640 with FBS and penicillin-streptomycin at 37°C and 5% CO2. mESCs were maintained in 2i medium and dissociated with Accutase for nuclei isolation. The PDX model GBM39 was cultured in DMEM/F12 with growth factors and supplements, including continuous erlotinib for GBM39-ERL. Adherent cells were washed, trypsinized, and centrifuged for collection, while suspension cells were collected directly by centrifugation. Cell pellets were resuspended in PBS for crosslinking.

### Nuclei preparation from the mouse cortex

All animal work described in this manuscript has been approved and conducted under the oversight of the Institutional Animal Care and Use Committee at the University of California, San Diego. Male C57BL/6J mice were purchased from the Jackson Laboratory (000664) at 7 weeks of age and were housed in the animal facility at University of California, San Diego, under a 12-h light/12-h dark cycle in a temperature-controlled room with *ad libitum* access to water and food until euthanasia and tissue collection at 8 weeks of age. The isocortex was dissected from 8-week-old male mice, snap-frozen in liquid nitrogen and stored at −80 °C before proceeding to nuclei extraction.

Single-nuclei suspensions were prepared from fresh tissues using dounce homogenization in a buffer containing sucrose, KCl, MgCl2, Tris-HCl, DTT, protease inhibitors, RNaseOUT, SUPERaseIn inhibitor, and Triton-X100. The suspension was filtered through a 30-μm filter and centrifuged at 300 x g for 10 minutes at 4°C. Nuclei were washed with buffer without Triton-X100, centrifuged, and resuspended in PBS at a concentration of 1 million cells/mL for crosslinking.

### Droplet Hi-C procedure

Cells were fixed with 1% formaldehyde, quenched with glycine, washed, and flash frozen at −80°C. For in situ Hi-C, cells were lysed in cold lysis buffer, treated with SDS and Triton X-100, and digested with restriction enzymes DpnII, MboII, and NlaIII. After deactivating enzymes, cells underwent a ligation reaction. For single-cell Hi-C sequencing library construction, the cell pellets were resuspended in 1 mL of PBS with 1% BSA. Cells were stained with 7-AAD and sorted for single nuclei using a fluorescence-activated nuclei sorter. Nuclei were collected, centrifuged, and resuspended in 1x Nuclei Buffer. Following cell counting with DAPI staining, nuclei were aliquoted for tagmentation using Chromium Next GEM Single Cell ATAC Reagent Kits v1.1 or v2. This step included an extended index PCR elongation time and a double-sided size selection process using SPRIselect beads to selectively remove smaller DNA fragments. Sequencing was completed on an Illumina NextSeq 2000 or NovaSeq 6000.

### Paired Hi-C procedure

Cells were crosslinked using 0.6% formaldehyde diluted in PBS, incubated for 10 minutes at room temperature, and quenched with glycine. Following centrifugation at 1,000g, cells were washed in PBS with BSA, pelleted, and flash-frozen at −80°C. For in situ Hi-C, lysed cell pellets underwent enzymatic digestion with DpnII, MboII, and NlaIII in a buffer containing Tris-HCl, NaCl, Igepal, and RNase inhibitors. The mixture was then treated with SDS, quenched, and ligated using T4 DNA ligase in a ligation buffer.

For single-cell sequencing library preparation, ligated cells were resuspended in PBS with BSA and RNase inhibitors, stained with 7-AAD, and sorted for single nuclei isolation. Nuclei were processed using Chromium Multiome kit (ATAC + Gene Expression analysis), followed by library preparation involving reverse transcription, barcoding, and size selection using SPRIselect. Sequencing was performed on an Illumina NextSeq 2000 or NovaSeq 6000 sequencer.

### Processing of Droplet Hi-C data

Custom scripts for demultiplexing, mapping and extracting single cell contacts from Droplet Hi-C data were developed. First, depending on indexing primers used, Droplet Hi-C fastq files were demultiplexed using Illumina bcl2fastq (v2.19.0.316) or 10X Genomics cellranger-atac mkfastq (v2.0.0). After demultiplexing, a custom script was used to extract cellular barcode sequence from Read2, and barcodes are aligned to the whitelist provided by 10x Genomics using Bowtie (v1.3.0)^70^. Aligned barcode sequences were appended to the beginning of the read name to record cellular identity of each read. Next, sequencing adapters were detected and trimmed with Trim-Garole (v0.6.10)^71^. Cleaned reads were then mapped to the human (hg38) or mouse (mm10) reference genome using BWA-MEM (v0.7.17)^72^, with arguments ‘- SP5M’ specified. Finally, after mapping, valid contacts were parsed, sorted and deduplicated by Pairtools (v0.3.0)^73^, with barcodes information stored in a separated column in the pairs file. Contacts were balanced and stored in cool format using Cooler (v0.8.10) for visualization and downstream analysis^74^. High-quality cells were selected based on the number of total unique contacts per cell in each library.

### Analysis of Droplet Hi-C data

We here described the general analysis strategy and workflows for our Droplet Hi-C data. Dataset-specific manipulations in the manuscript, if any, will be indicated.

#### Visualization of Hi-C contact matrices

Bulk, pseudo-bulk, or single cell contact matrices in cool format were visualized and plotted using Cooler along with Matplotlib (v3.5.1)^75^.

#### Embedding and clustering of single-cell Droplet Hi-C dataset on cultured cells

We used Higashi to infer low-dimensional cell embeddings for all our Droplet Hi-C datasets without imputations, including human cell lines mix (HeLa / GM12878 / K562, Extended Data Fig.1a), GBM39 / GBM39-ER (Fig. 4b), and GBM patient sample (Fig. 5b). For visualization, the L2 norm of cell embeddings were projected to two-dimensional space with uniform manifold approximation and projection (UMAP). To identify cells with similar identity, we performed Leiden clustering using igraph (v0.9.9) and leidenalg (v0.8.8)^76, 77^.

#### Imputation of single-cell chromatin contacts on adult mouse cortex Droplet Hi-C dataset

For the adult mouse cortex Droplet Hi-C data, we used scHiCluster (v1.3.4) to perform contact matrix imputation for individual cell at three different resolutions: 100 kb (for cells embedding and visualization), 25 kb (for domain boundaries calling) and 10 kb (for cells embedding and loops calling)^78^. In brief, contacts from individual cell were first masked for ENCODE blacklist regions^79^. scHiCluster then performed linear convolution and random walk with restart to impute the sparse single cell contact matrices for each chromosome. Considering storage efficiency, we output the imputed results for the whole chromosome at 100kb, for 10.05Mb window at 25kb, and for 5.05Mb window at 10kb resolution for each cell.

#### Analysis of A/B compartments

On sample levels, we used the ‘eigs_cis’ module from cooltools (v0.5.1) to perform eigen decomposition on balanced contact matrices and calculate the compartment score at 100 kb resolution^80^. The orientation of the resulting eigenvectors was adjusted to correlate positively with the CpG density of the corresponding 100 kb genomic bins, thereby determining the sign of the compartments (A/B). Similarity between samples is calculated as the Spearman correlation coefficient of compartment scores across all autosomes (Extended Data Fig. 1c). To illustrate chromatin interactions within and between compartments, we used the ‘saddle’ module from cooltools to calculate the average observed versus expected contact frequency, categorized by compartment scores.

To generate pseudo-bulk profiles for distinct cell types or clusters within a sample, we adopted a uniform approach outlined before to minimize bias and facilitate direct comparisons. Initially, the raw contact matrices at the sample level were normalized for distance effects. Next, Pearson correlation coefficients were computed on the distance-normalized matrices, and PCA was then performed on the correlation matrix. The model fitted at the sample level was utilized to transform the raw contact matrices of various cell subtypes or clusters. To compare similarity between cell types or clusters, we calculated the Spearman correlation coefficient of compartments score using compartment intersected with variable gene regions (Fig. 1e, Extended Data Fig. 7c). To identify differential compartments, we employed dchic (v2.1)^81^, selecting only those with an adjusted P value < 0.01 and a Manhattan distance greater than the 2.5^th^ percentile of the standard normal distribution for further downstream analysis.

For joint analysis of compartment and cell type-specific histone modifications, we utilized the recently published adult mouse cortex Droplet Paired-Tag dataset^38^. We retained only cell types that contain >100 cells, and were shared between datasets to ensure comparability across the analyses. For all differential compartments, we calculated the Pearson correlation coefficients between compartment score and signal of H3K27ac or H3K27me3 (CPM) within the same 100 kb genomic bins across various cell types.

#### Analysis of chromatin domains

On sample levels, we used the “insulation” module from cooltools to compute insulation score for raw contact matrices at 25 kb resolution. To delineate cell type-specific variable domain boundaries in mouse cortex data, we employed TopDom (v0.0.2) on 25 kb resolution imputed contact matrices for each individual cell^82^. We defined the ‘boundary probability’ for a given bin as the proportion of cells within a particular cell type that designate the bin as a domain boundary. The presence or absence of a domain boundary is summarized by n cell types into a n × 2 contingency tables. We then computed the chi-square statistics and P value for each bin, and the domain boundaries were classified as variable if they exhibited a false discovery rate (FDR) < 0.001 and a boundary probability difference > 0.05.

For correlation analysis of gene expression surrounding variable domain boundaries, we first identified genes within a 100kb window centered on the variable domain boundaries. We then calculated Pearson correlation coefficients to quantify the relationship between boundary probabilities and expression levels (RPKM) of these genes from Droplet Paired-Tag data across shared cell types. The genes were further categorized into “housekeeping”, “constant”, and “variable”. In brief, the housekeeping gene list is taken from previous report^83^. The top 2,500 variable genes from Droplet Paired-Tag RNA modality are selected as variable group. Genes with similar expression level as the top 2,500 variable genes but are not within the “housekeeping” or “variable” group is treated as “constant”.

#### Analysis of chromatin loops

For loops calling on sample levels, we adopted the “dots” module in cooltools, which implements the principle of the commonly used HiCCUPS loop calling strategy, to call loops on the 10 kb raw contact matrices^84^.

A modified version of SnapHiC workflow implemented in scHiCluster is used to perform loops calling on the 10-kb imputed contact matrices for mouse cortex cell types^85^. To compare histone modification enrichment at loop anchors, we calculate H3K27ac or H3K27me3 CPM across all cell types at all loop anchors identified. When histone modification profiles and loop anchors are from the same cell types, the enrichment is classified as “matched”, otherwise classified as “unmatched”.

Gene Ontology annotation of loop anchors was performed using rGREAT (v1.26.0) in R^86^. Gene Ontology biological process was selected for annotations. The result is used to generate plots in Fig. 2g

#### Infer copy number variation (CNV) with Droplet Hi-C data

Copy number is inferred from Hi-C data using the ‘calculate-cnv’ module in NeoLoopFinder (v0.4.3)^87^. For each single cell, the output residuals from the generalized additive model in ‘calculate-cnv’ was directly used to estimate copy number ratio. On bulk or pseudo-bulk level, an additional HMM segmentation is performed using the ‘segment-cnv’ module on the copy number ratio to determine the boundaries of copy number ratio segments. We assume all samples used in this manuscript for CNV calculation are diploid, therefore the inferred copy number is equal to 2 × copy number ratio. Genome-wide CNV heatmap is plotted at 10 Mb resolution, where chromosome level CNV is plotted at 1 Mb resolution, and regional CNV profile is plotted at 10 kb resolution in R using pheatmap (v1.0.12). The inferred copy number is further used to assist identifying ecDNA candidates and the associated significant *trans-* interactions, and to correct the bias in contact matrices for structural variation identification.

#### Identify structural variation (SV) from Droplet Hi-C data

Structural variations (SV) were identified using the ‘predictSV’ module in EagleC (v0.1.9)^88^. In short, CNV effects on contact matrices at 5, 10, and 50 kb resolutions were removed using ‘correct-cnv’ module in NeoLoopFinder. ‘predictSV’ then uses a deep learning model to predict SV at each resolution on the corrected matrices, and combines all results to obtain a uniform, high-resolution SV list at 5 kb resolution. ‘annotate-gene-fusion’ module is applied on the final SV list to annotate gene fusion events.

#### Metrics to define chromatin interactions

We compiled a suite of metrics to assess the *cis-* and *trans-*interaction patterns at specific genomic loci. These metrics include: (1) contact evenness or “hub index”, quantified as the Gini coefficient for *trans-*interactions across all chromosomes excluding chromosome Y given an 1 Mb genomic bin, and was derived using ineq (v0.2-13) in R; (2) *trans-to-cis* contacting bin ratio (*R_i_*), which compares the quantity of interacting bins within the same chromosome (*N_C_*) to those across different chromosomes (*N_T_*) for a given bin *i*: 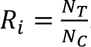. This ratio represents the *trans-*interaction tendency while it is not confounded by copy number variation; (3) copy number-adjusted *trans-*chromosomal interaction (*adj_n_TIF*), which is also used to measure a genomic locus’s interaction activity, was calculated as described before^44^.

### Develop ecDNA caller for identifying ecDNA candidates

We developed two ecDNA identification methods using regression and convolutional neural network (CNN) models to predict ecDNA presence in 1 Mb genome bins. The regression model employed a logistic regression trained on COLO320DM and GBM39 datasets with defined ecDNA loci as positive examples, using copy number, hub index, and bin contact ratios as predictors. A specific probability threshold was established for each cell line to differentiate ecDNA presence, set at 0.5 for COLO320DM/COLO320HSR and 0.95 for GBM39/GBM39-ER.

The CNN-based model was trained using Hi-C contact matrices binarized at 1 Mb resolution, incorporating both cis- and trans-contacts within a 7 Mb neighborhood as input features. This architecture included two convolutional modules with sequential batch normalization, ReLU activation, and max-pooling layers, followed by a fully connected network with a dropout layer to enhance model robustness and prevent overfitting. The model, trained over 40 epochs with the AdamW optimizer and a learning rate scheduler, utilized bootstrapped cross-entropy loss, biased class weights, and evaluated performance using precision, recall, and accuracy metrics. Final ecDNA predictions per cell were determined using SoftMax probabilities, with an aggregate analysis to estimate ecDNA and HSR prevalence in the cell population.

### Identify significant *trans-*interactions of ecDNA candidate loci

To identify genomic regions preferentially interact with ecDNA, we first quantify the number of interactions (*N_i_*) of 500 kb genomic interval *i* on different chromosomes with ecDNA candidate loci, treating each chromosome separately. The observed interaction frequency (*P_i_*) is the proportion of *N_i_* relative to all contacts on the same chromosome: 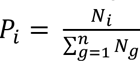 The expected interaction frequency for interval *i* was calculated as the ratio of the inferred copy number (*CN_i_*) to the total copy number on the same chromosome: 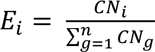 based on the null hypothesis that interaction frequency for genomic regions with ecDNA are only weighted by their underlying copy number. To identify significant interactions, observed-versus-expected *P* value was calculated based binomial distribution model. Multiple testing correction is done by Bonferroni adjustment. Regions with adjusted *P* value < 0.05 are selected as significant interacting regions.

### Analysis of 10X Multiome datasets

10X Multiome fastq files for the GBM patient sample were demultiplexed and pre-processed with cellranger-arc (v2.0.0). Clustering of single-nucleus transcriptomic or chromatin accessibility data were performed in R using Seurat (v4.1.0) or Signac (v1.6.0)^89, 90^. For transcriptomic data, gene counts were normalized and scaled, and the top 2,500 variable genes were selected for dimension reduction by PCA. The first 30 principal components were used for UMAP visualization and Louvain clustering. Potential doublets were identified and removed using Scrublet^91^. For chromatin accessibility data, cell-by-peak matrices were normalized by the two-step term frequency-inverse document frequency (TF-IDF). The top 95% of genomic bins were selected for linear dimension reduction, again followed by UMAP visualization and Louvain clustering. Gene activity scores were computed by ATAC signal density in promoter and gene body regions.

### Analysis of GBM cellular states

To classify single cells within GBM 10x Multiome or Paired Hi-C RNA datasets into one of four predefined malignant cellular states, we adopted a previously described two-dimensional visualization technique^49^. Briefly, we quantified the gene enrichment score (*SC_j(i)_*) for each cell *(i)* against one of the four gene sets (*G_j_*) associated with the particular cellular state. This score was calculated as the relative averaged expression (*Exp*) of *G_j_* in cell *i* compared to a group of genes (*G*) with a similar level of expression as control: 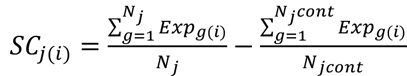 where *N_j_* and *N_jcont_* are the number of genes in *G_j_* and *G_jcont_*. After obtaining scores for all four cellular states, the cells were stratified into OPC/NPC versus AC/MES categories using the differential 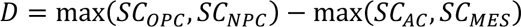. For further refinement, OPC/NPC cells were assigned an identity value 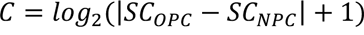, and AC/MES cells were similarly categorized with 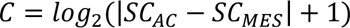. The distribution of cellular states was then plotted in the two-dimensional representation with *D* on the y-axis and *C* on the x-axis.

For cellular state identification in Droplet Hi-C data, co-embedding with the reference snRNA-seq was performed using the scGAD score. After determining the top 15 nearest snRNA-seq neighbors and their scaled distance-based similarity score (*D_m_*) for each Droplet Hi-C cell, the Hi-C gene enrichment score (*HSC_j(i)_*) was computed as the sum of the neighbor-weighted enrichment scores: 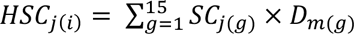.

### Single-cell genotyping by Droplet Hi-C data

Frequently observed mutations in AML were sequenced using Hematologic Malignancy Comprehensive Panel for the patient BMMC sample. To detect malignant cells carrying the detected mutations from Droplet Hi-C data, cellular barcode information was appended to the mapped bam files as an extra tag. To ensure comparability between pre- and post-treated samples, bam files were subsampled to a uniform sequencing depth. we employed a previously reported custom script to count reads for both the wild-type and mutant alleles, considering only barcodes that met predefined quality standards^92^. For each cell and mutational site, we then summarized the detected mutant and wild-type reads. Given the limited sequencing coverage, we defined potential malignant cells as cell carrying at least one mutant read.

### Processing and Analysis of Paired Hi-C data

Preprocessing and analysis of Hi-C modality in Paired Hi-C are identical as described for Droplet Hi-C. For RNA modality, fastq were demultiplexed by cellranger-arc, but preprocessed using cellranger (v6.1.2). After obtaining the cell-by-gene matrix, clustering and visualization were performed as described for 10x Multiome RNA dataset. Since barcodes in the same gel bead for RNA and Hi-C modalities are different, we performed manual pairing to match cell barcodes based on the 10X Multiome barcodes whitelist provided in cellranger-arc (10x Genomics).

Integration of the Paired Hi-C RNA dataset with reference datasets was performed by Seurat. First, normalization of gene counts was performed, and the top 2,000 shared variable genes across datasets were selected as integration features. Subsequent canonical correlation analysis allowed projecting all nuclei into a unified embedding space. Anchors (pairs of corresponding cells from distinct datasets) were then discerned through mutual nearest neighbors searching. Anchors with low confidence were excluded, and the shared neighbor between anchors and query cells are computed. Louvain clustering was applied to the shared neighbor graph to discern co-embedded clusters. We calculated overlap coefficients as described in previous section to compare clustering results from different datasets.

### Analysis of gene expression levels and copy numbers of ecDNA

To calculate the correlation between gene expression level and inferred copy number of genes on ecDNA, we first refined the range of ecDNA at 10 kb resolution. This is based on the observation of the increased intra-ecDNA interaction frequency than interactions with regions on linear genome, irrespective of their genomic distance. Specifically, we enumerated the local interactions within each 10 kb segment of the 1 Mb ecDNA candidates. Subsequently, the contact numbers were smoothed across adjacent bins. By calculating the differential contact numbers between neighboring bins, we determined the changing points of interaction with an average difference cutoff across the entire 1 Mb region. The outermost local maxima and minima were designated as ecDNA boundaries.

With the ecDNA boundaries established, we categorized genes with gene bodies residing within ecDNA, as “ecDNA genes”. In the case of MYC ecDNA (ecMYC), we further classified genes on GBM39 ecMYC as “shared genes”, and genes on non-overlapping regions between GBM39 and GBM39-ER as “ecMYC variable genes”, considering the greater ecDNA size in GBM39-ER at pseudo-bulk level. The Spearman correlation coefficient between gene expression levels (RPKM) and the average inferred copy number over all 10 kb segments at gene body across all cells was calculated, for a comprehensive representation of the interplay between gene expression and copy number.

### Ethical approval

The GBM specimen collection was approved by the Institutional Review Board (IRB) at the University of Minnesota. The AML and MDS specimen collection was approved by the IRB at the University of California, San Diego. Each patient was consented by a dedicated research coordinator prior to collection. Samples were obtained with informed consent in accordance with the Declaration of Helsinki and appropriate Ethics Committee approval from each partner institution.

## Acknowledgements

We are grateful to Dr. Jingtian Zhou for providing suggestions in data analysis. We thank Drs. Jie Xu and Chenxu Zhu for their invaluable suggestions on this study. The work was supported by NIH grants 1UM1HG011585 (to B.R. and M.H.), R35HG011922 (to M.H.), R56 NS080939 and R01 CA258248 (to F.B.F.), NIH training grants (5T32CA067754-27 and 5T32GM007198-47 to B.T.). Z.W. is a DDBrown awardee of the Life Sciences Research Foundation. This publication includes data generated at the UC San Diego IGM Genomics Center utilizing an Illumina NovaSeq 6000 that was purchased with funding from a National Institutes of Health SIG grant (#S10 OD026929)

## Author information

### Contributions

This study was conceived by B.R. and L.C. L.C. designed and implemented all the Droplet Hi-C, Paired Hi-C and part of 10x Multiome experiments. Y.X. performed all the bioinformatics analysis. B.T. contributed to ecDNA related experimentation and data analysis. Z.W. dissected mouse cortex samples for this study, generated 10x multiome data from GBM samples, and performed next-generation sequencing on the Nextseq 2000 platform. M.H. and J.S. developed ecDNA caller algorithm. R.B. and T.T. provided the AML patient samples, C.C. provided the GBM patient samples and F.F. provided the GBM39 cell line and related reagents. B.R., L.C., and Y.X. wrote the manuscript with input from all the authors.

### Corresponding author

Correspondence to Bing Ren.

---

**Extended Data Fig. 1.**
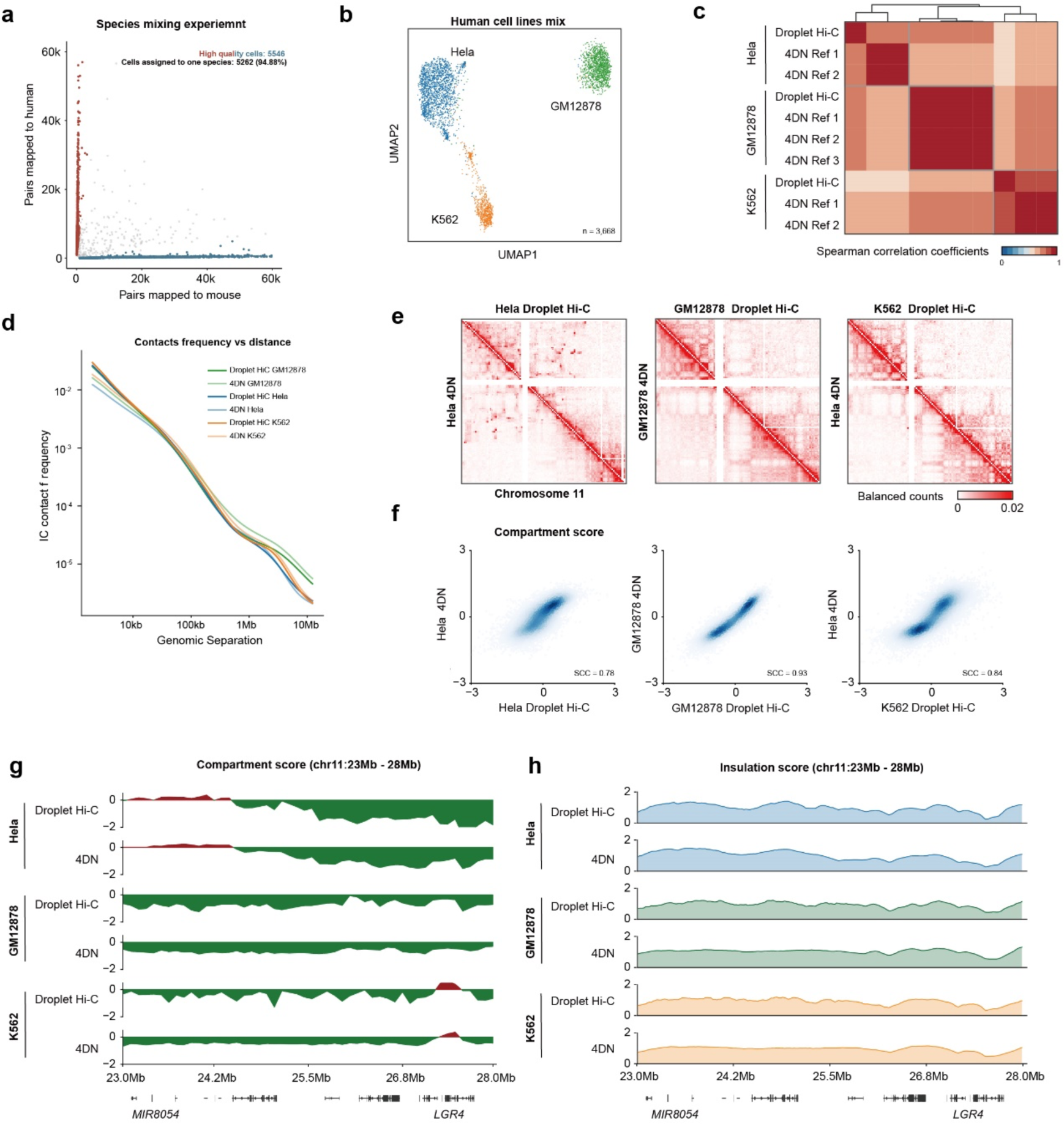
Comparison between Droplet Hi-C and *in situ* Hi-C on cultured cells. **a,** Scatter plots showing proportion of human and mouse DNA reads in each cell in the Droplet Hi-C species mixing experiment. **b,** UMAP visualization of Droplet Hi-C data on three human cell lines mixing sample. **c,** Genome-wide Spearman correlation coefficients (SCC) of compartment score between pseudo-bulk Droplet Hi-C data and bulk *in-situ* Hi-C datasets. **d,** Comparison of contact frequency by distance between Droplet Hi-C and *in situ* Hi-C datasets of Hela S3, GM12878 and K562 cells. **e,** Comparison of contact maps between Droplet Hi-C and *in situ* Hi-C for Hela S3, GM12878 and K562 cells on chromosome 11 at 100-kb resolution. **f,** Scatter plot showing genome-wide SCC of compartment score between Droplet Hi-C and *in situ* Hi-C for Hela S3, GM12878 and K562 cells. **g-h,** Comparison of compartment score **(g)** and insulation score **(h)** profiles on chromosome 11: 23 - 28 Mb from Droplet Hi-C and *in situ* Hi-C datasets for Hela S3, GM12878 and K562 cells.

**Extended Data Fig. 2.**
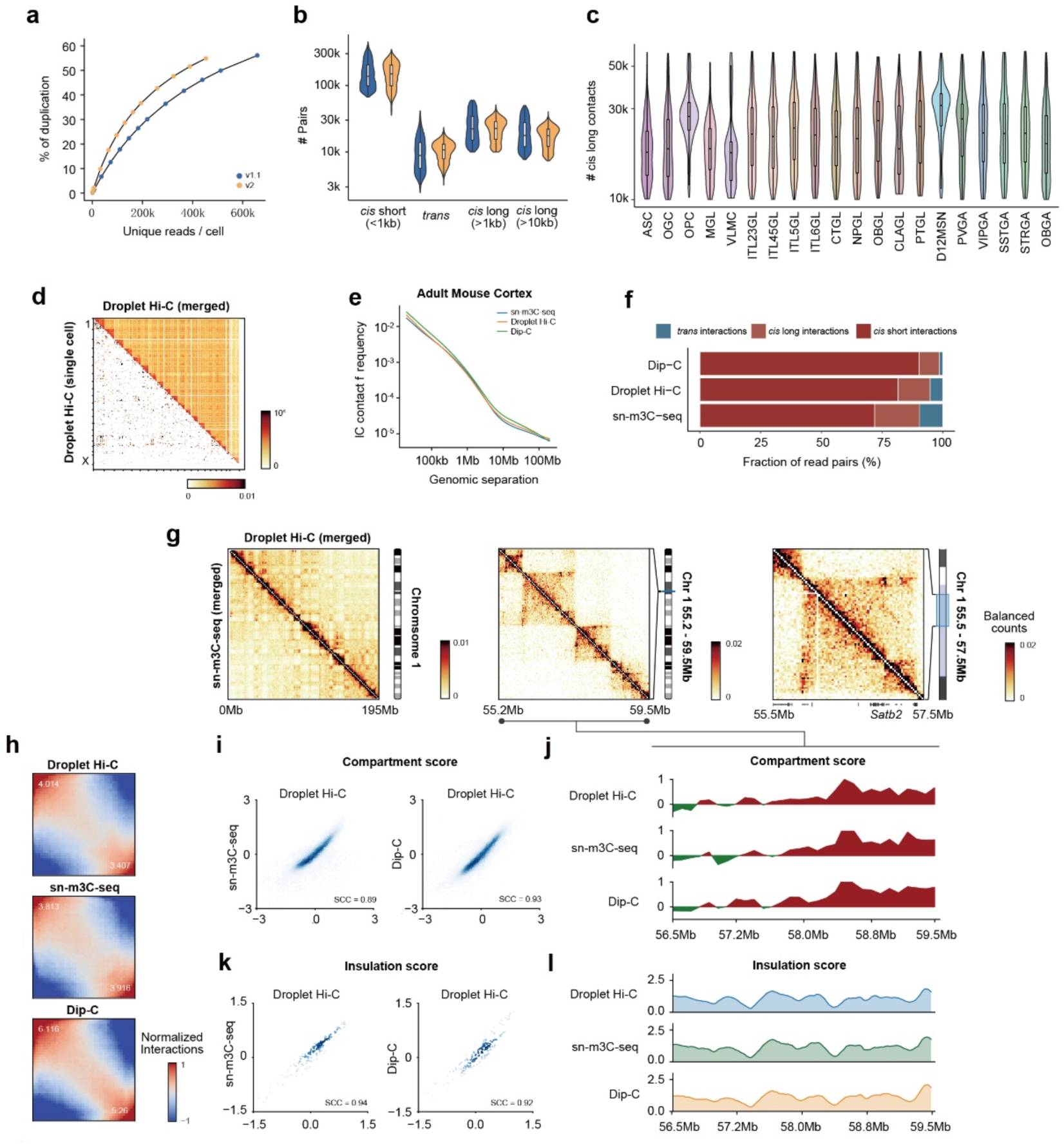
Performance of Droplet Hi-C on adult mouse cortex. **a,** Comparison of library complexity between two datasets generated with 10x Genomics Chromium Single Cell ATAC kit v1.1 and v2. **b,** Distribution of *cis*-short*, cis*-long or *trans-* contacts number per nucleus. **c,** Comparison of *cis*-long contact (>1kb) numbers distribution across different cell types. **d,** Pseudo-bulk and representative single-cell genome-wide contact maps from mouse cortex. Color bar showed the raw contact number. **e-f,** Comparison of contact frequency by distance **(e)** and contacts ratio **(f)** on mouse cortex data among Droplet Hi-C, sn-m3C-seq and Dip-C. The *cis*-long interactions in **(f)** represent contacts separated more than 1 kb. **g,** Comparison of multi-scale genome organization on chromosome 1 between Droplet Hi-C and sn-m3C-seq. **h-l,** The compartmentalization strength **(h)**, compartment scores **(i-j)** and insulation scores **(k-l)** among Droplet Hi-C, sn-m3C-seq and Dip-C are also shown.

**Extended Data Fig. 3.**
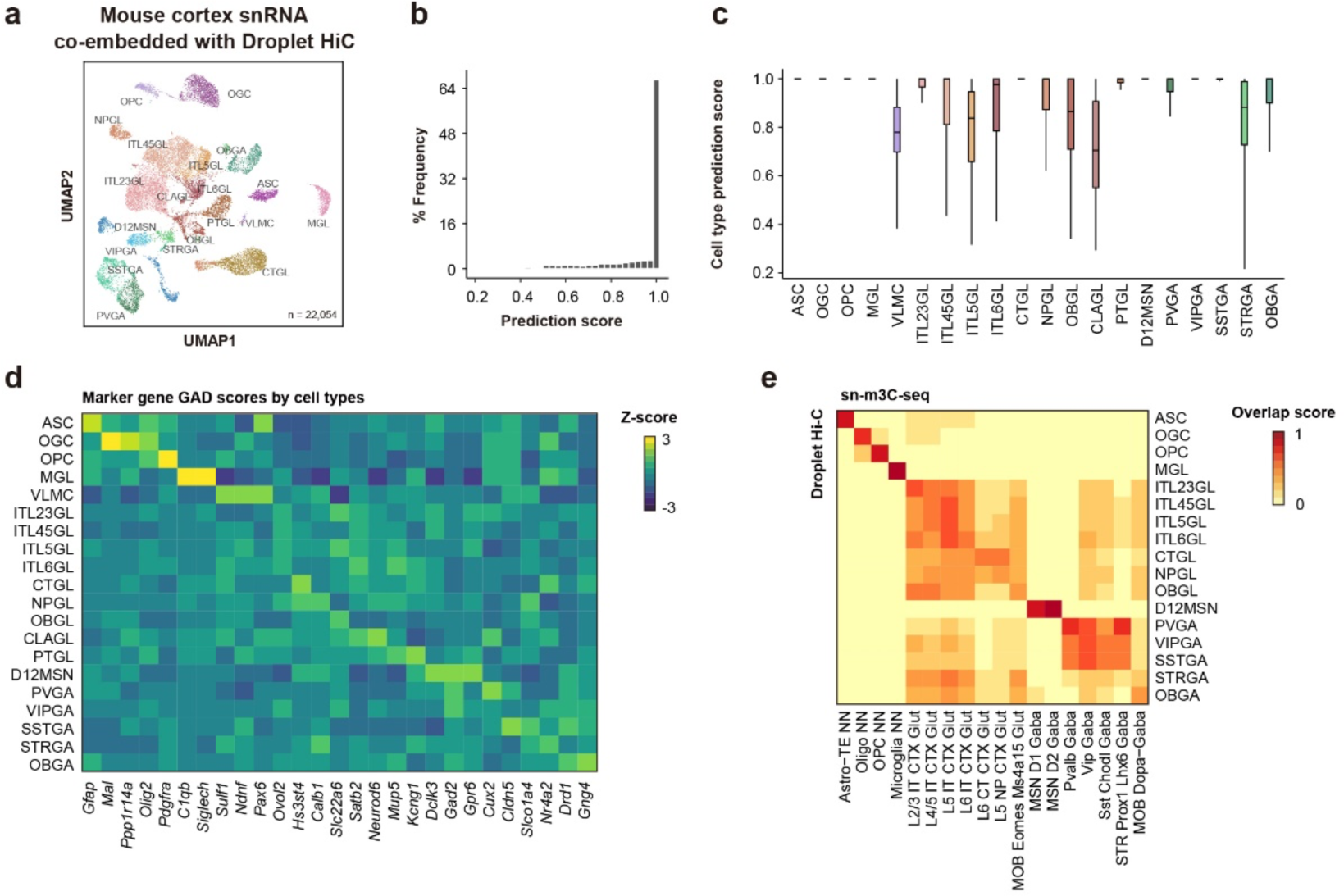
Validation of cell type identities in mouse cortex characterized by Droplet Hi-C. **a,** UMAP visualization of public mouse cortex single-nucleus RNA-seq data used to co-embed Droplet Hi-C. **b,** Distribution of single-cell prediction scores for Droplet Hi-C cells, embedded with Droplet Paired-Tag RNA data. **c,** Comparison of single-cell prediction scores across different cell types. **d,** Heatmap showing gene associating domain (GAD) scores for known marker genes among all the cell types from Droplet Hi-C data. **e,** The overlap scores between the joint clusters and the original annotations from Droplet Hi-C and sn-m3C-seq data in adult mouse cortex.

**Extended Data Fig. 4.**
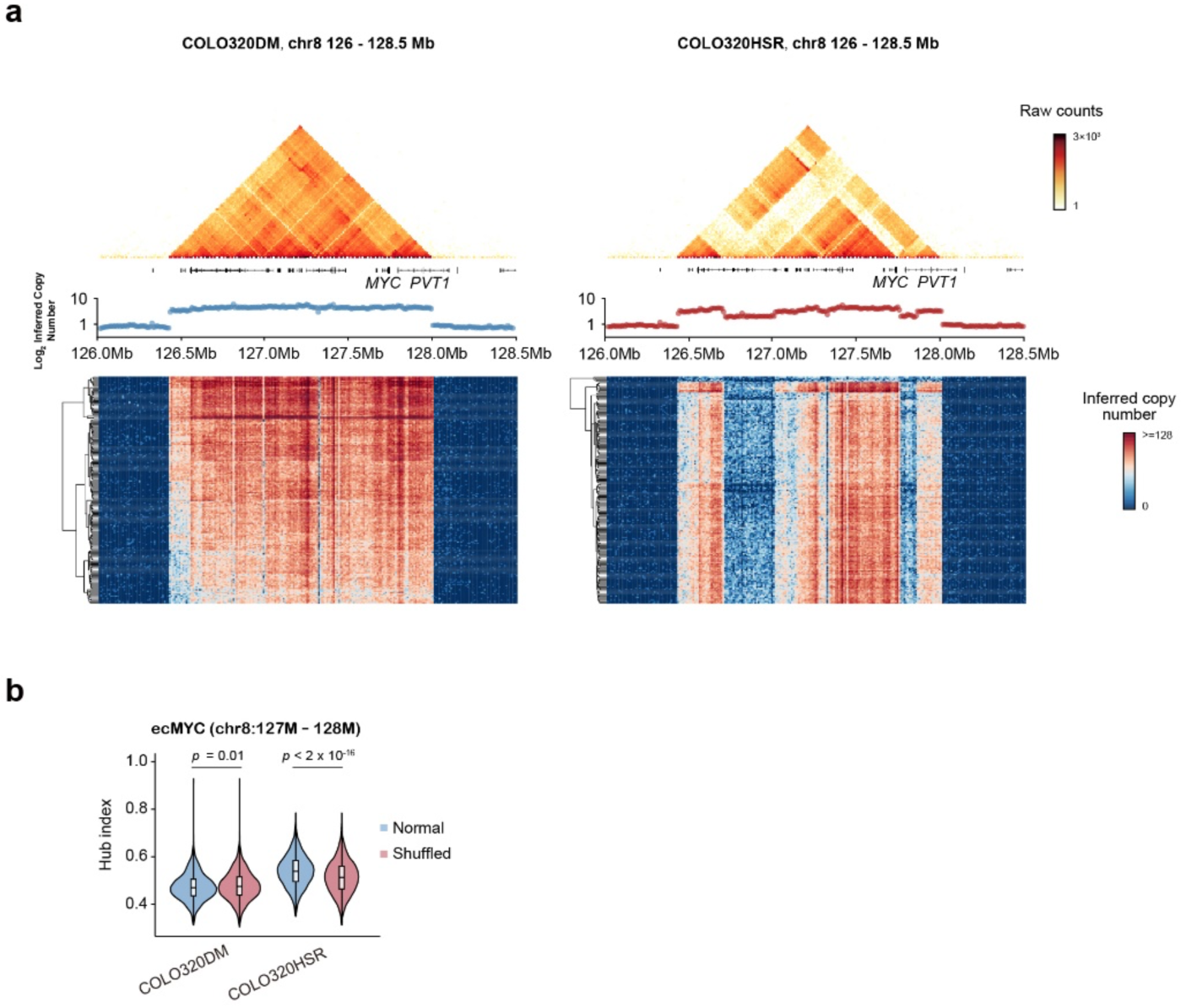
Droplet Hi-C illuminates CNV, SVs and ecDNA in cancer cells. **a,** Contact maps of regions containing ecMYC in COLO320DM and COLO320HSR. Inferred copy number from pseudo-bulk profiles as well as representative single cells are shown below. **b,** Single-cell hub index calculated with normal versus shuffle contact profiles in COLO320DM and COLO320HSR. *P* values are calculated from paired sample Wilcoxon signed-rank test.

**Extended Data Fig. 5.**
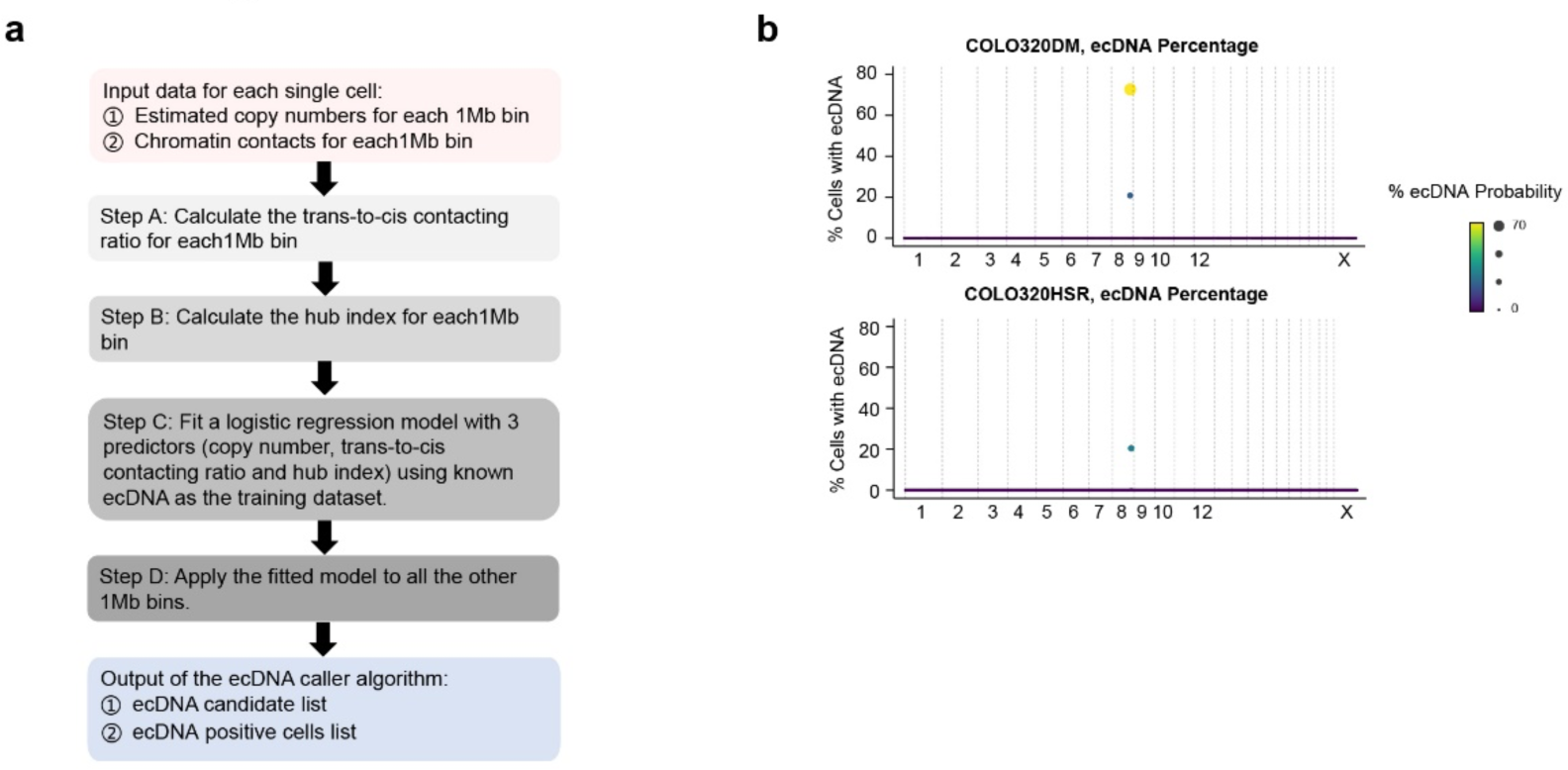
Flowchart of ecDNA caller algorithms. **a,** Workflow of multivariate logistic regression model-based ecDNA caller used to predict ecDNA identity for each genomic region, and cells containing the candidate ecDNA. **b,** ecDNA prediction results from the logistic regression model-based ecDNA caller in COLO320DM and COLO320HSR sample.

**Extended Data Fig. 6.**
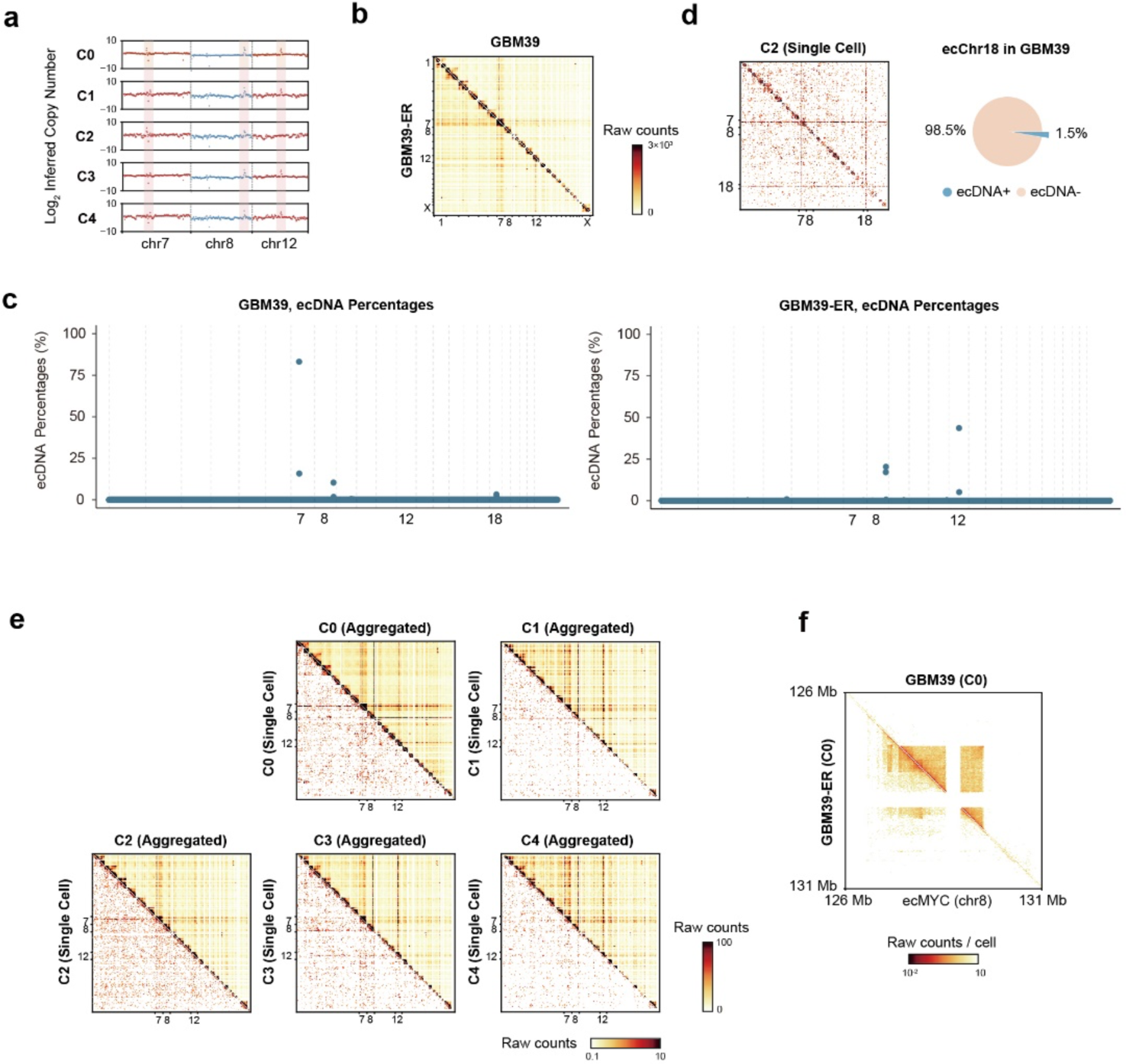
Dynamics of ecDNA in GBM39 under erlotinib treatment. **a,** Scatter plot showing inferred copy number on chromosomes containing candidate ecDNA (chromosome 7, 8, 12) across clusters in GBM39 and GBM39-ER. ecDNA-associated regions are highlighted in pink. **b,** Genome-wide contact maps for GBM39 and GBM39-ER cells. **c,** Genome-wide ecDNA prediction results from deep learning-based ecDNA caller for GBM39 and GBM39-ER. **d,** Representative single-cell contact map and percentage of cells containing rare ecDNA (ecChr18) found by deep learning-based ecDNA caller. **e,** Genome-wide contact maps at cluster and representative single-cell levels. **f,** Comparison of ecMYC interaction profiles for ecDNA positive cells within C0 from different samples.

**Extended Data Fig. 7.**
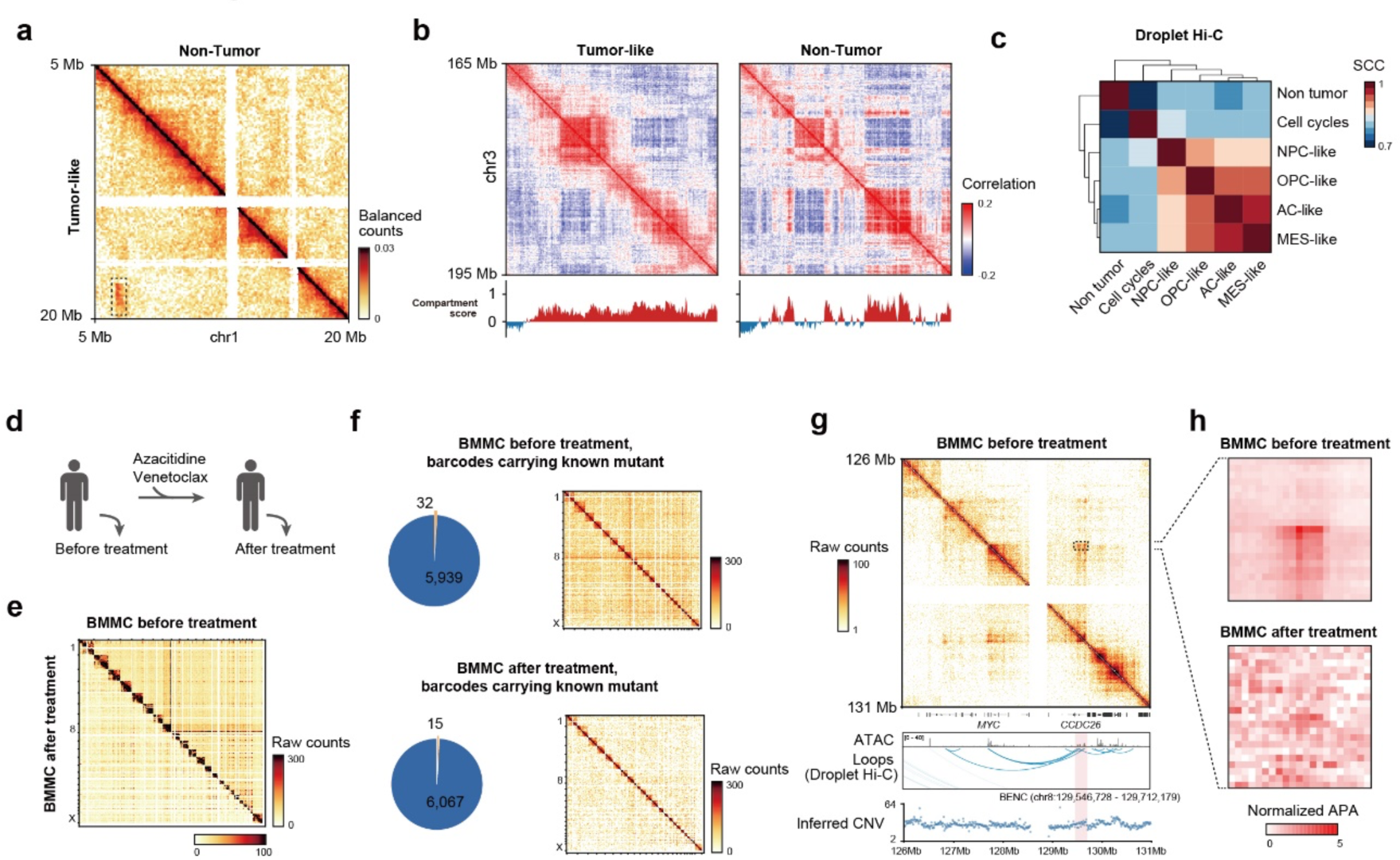
Cell state-specific chromatin structure and transcriptome differences in GBM patient samples. **a,** Another example of tumor-like population-specific SV on chromosome 1 from GBM patient sample. SV are highlighted with black rectangle. **b,** Comparison of A/B compartment profiles on chromosome 3 between tumor-like cells and non-tumor populations. **c,** Heatmap showing Spearman correlation coefficients (SCC) of Droplet Hi-C compartment score across different cellular states. **d,** Illustration of AML and MDS patient treatment procedure. **e,** Pseudo-bulk genome-wide contact maps for BMMC samples from AML and MDS patient before and after treatment. **f,** Proportion of cells containing known tumor-associated mutations in BMMC samples before and after treatment. Aggregated genome-wide contact maps from mutant-carrying cells are also shown. Color bar shows the raw contact number. **g-h,** An example of a before-treated sample-specific loop between *MYC* and its enhancer region (BENC) on ecMYC. Normalized aggregate peak analysis (APA) scores of before- and after-treated samples are shown aside.

**Extended Data Fig. 8.**
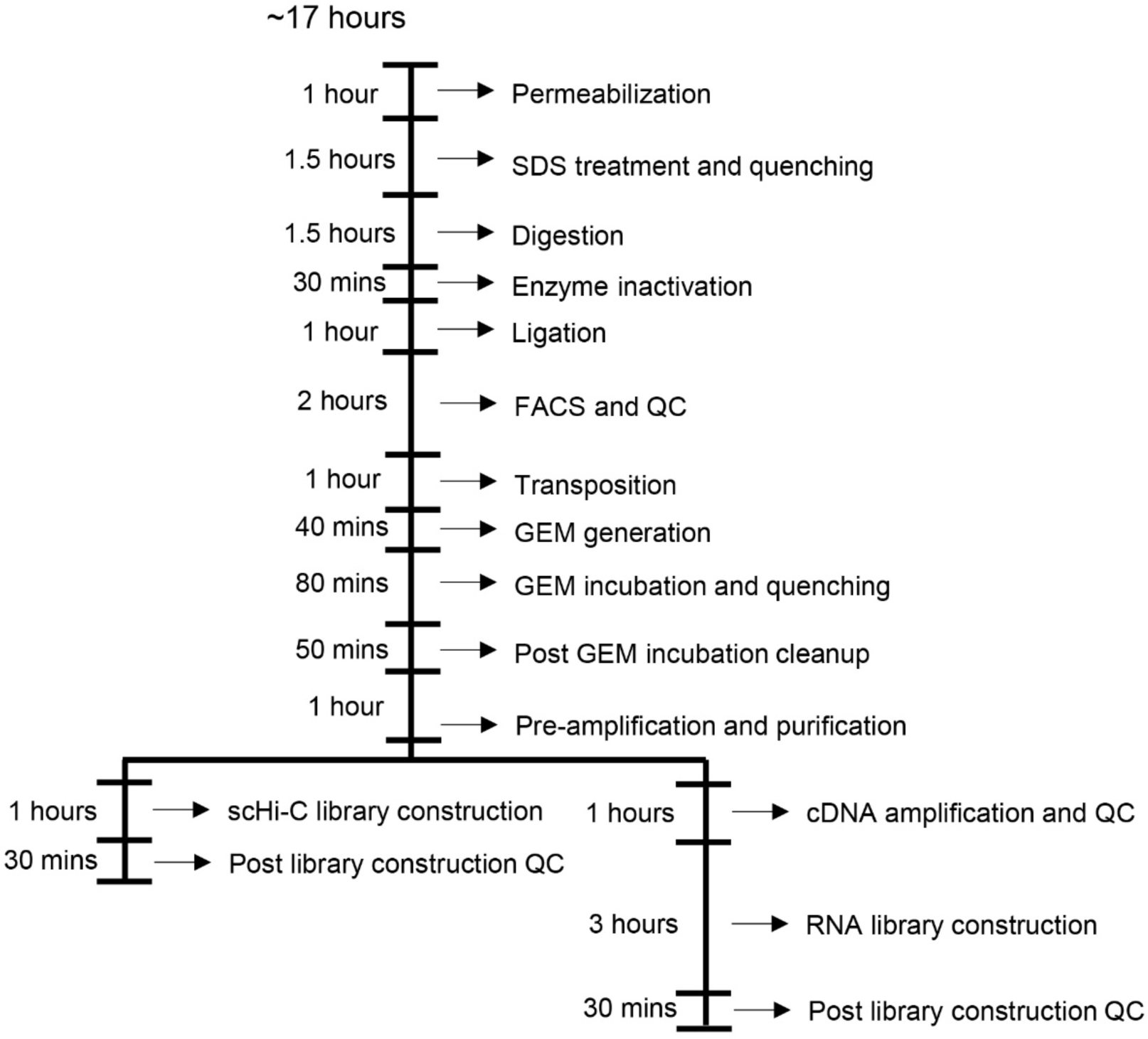
Workflow of Paired Hi-C. Detailed workflow describing the end-to-end procedure of Paired Hi-C and time cost for each step.

**Extended Data Fig. 9.**
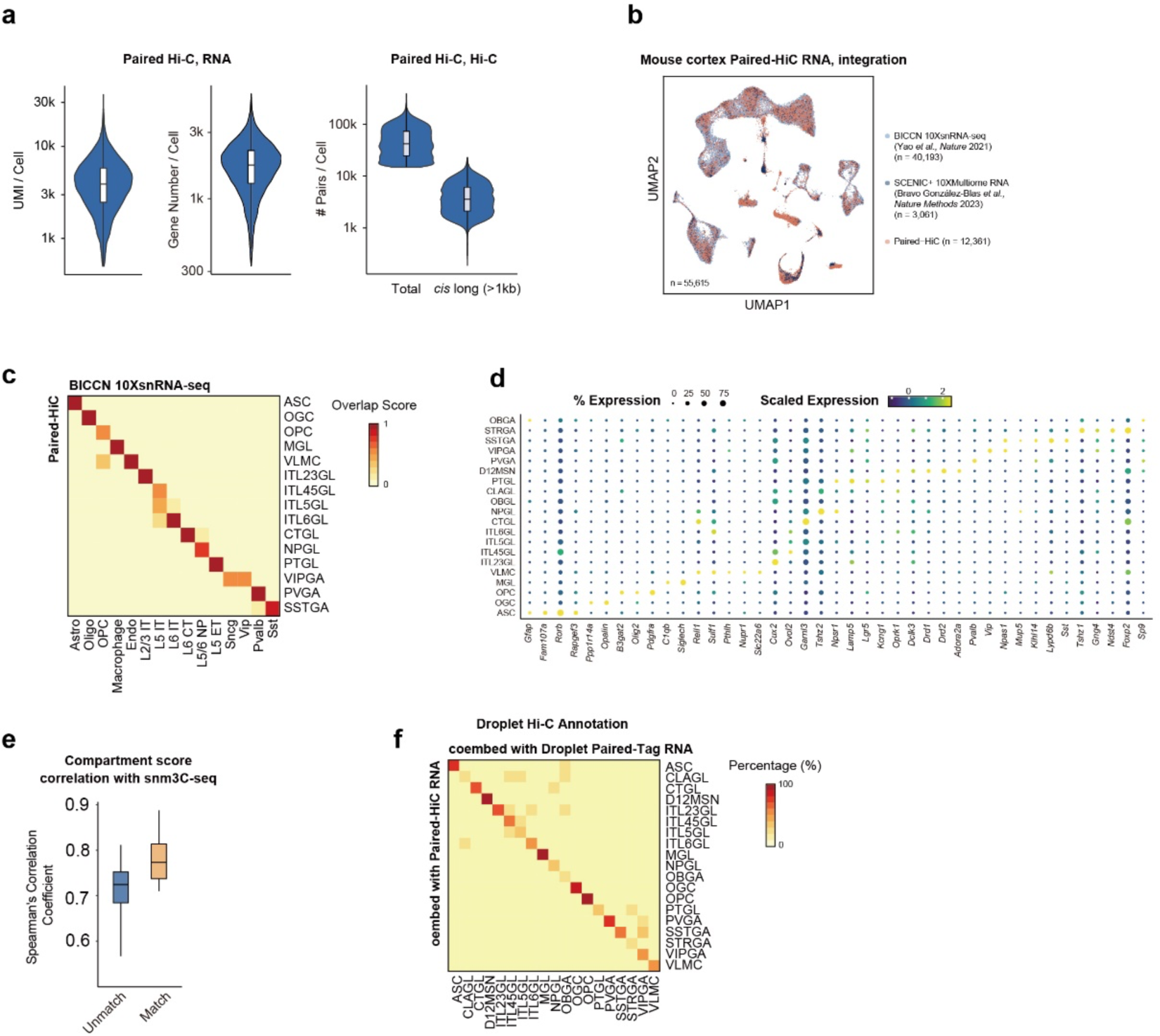
Joint chromatin organization and gene expression analysis using Paired Hi-C. **a,** Distribution of UMI, gene number, and contacts per nucleus in Paired Hi-C data from adult mouse cortex. **b,** UMAP co-embedding of single-nucleus transcriptome profiles from Paired Hi-C experiment and the reference BICCN and SCENIC+ datasets. **c,** The overlap scores of shared annotations between BICCN single-nucleus RNA-seq dataset and Paired Hi-C single-nucleus RNA-seq dataset. **d,** Dot plot showing expression level of marker genes in each cell type. **e,** Box plot showing Spearman correlation coefficients (SCC) of compartment score between sn-m3C-seq and Paired Hi-C. **f,** Overlap of co-embedding annotations between using Droplet Paired-Tag single-nucleus RNA-seq dataset as reference and Paired Hi-C single-nucleus RNA-seq dataset as reference.

**Extended Data Fig. 10.**
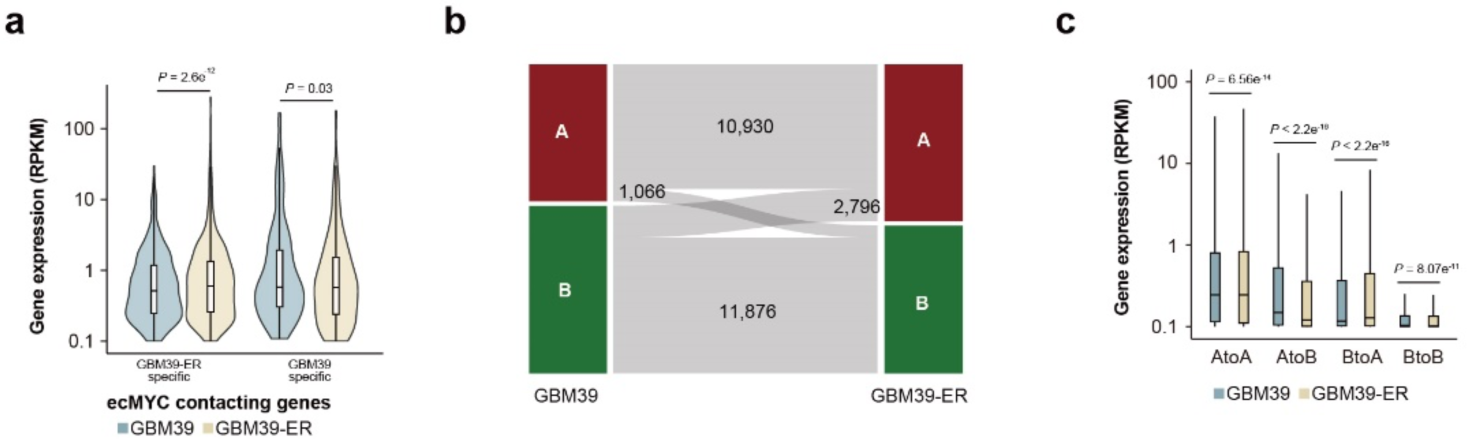
Paired Hi-C illuminates alterations in the transcriptome associated with variations in chromatin structure. **a,** Comparison of expression level for ecMYC *trans*-contacting genes between GBM39 and GBM39-ER cells. *P* values, Wilcoxon signed-rank test. **b,** Summary of changes in A/B compartments in GBM39 and GBM39-ER sample. **c,** Expression of genes classified by the underlying A/B compartment transitions in GBM39 and GBM39-ER cells.

